# Blm10/PA200-activated 20S proteasomes promote α-synuclein degradation and bypass proteasome inhibition in Parkinson’s disease models

**DOI:** 10.64898/2026.01.11.698857

**Authors:** Tariq T. Ali, Anton Zhornyak, Madiha Merghani, Zora Buschenlange, Eri Sakata, Tiago F. Outeiro, Blagovesta Popova, Gerhard H. Braus

**Affiliations:** Department of Molecular Microbiology and Genetics, Institute of Microbiology and Genetics, University of Göttingen, Grisebachstr. 8, 37077 Göttingen, Germany; University Medical Centre Göttingen, Department of Experimental Neurodegeneration, Centre for Biostructural Imaging of Neurodegeneration, Waldweg 33, 37073 Göttingen, Germany; Institute for Auditory Neuroscience, University Medical Centre Göttingen, Robert-Koch-Str. 40, 37075 Göttingen, Germany; German Centre for Neurodegenerative Diseases (DZNE), Von-Siebold-Str. 3a, 37075 Göttingen, Germany; Translational and Clinical Research Institute, Faculty of Medical Sciences, Newcastle University, Framlington Place, Newcastle Upon Tyne, NE2 4HH, UK

**Keywords:** alpha-synuclein, Parkinson disease, 20S proteasome, proteasomal chaperones, yeast, protein homeostasis, posttranslational modifications, autophagy

## Abstract

Protein homeostasis is essential for maintaining normal cellular function. However, protein homeostasis efficiency declines with age, leading to the accumulation of aberrant protein structures associated with neurodegenerative diseases such as Parkinson’s disease (PD). PD is characterized by the aggregation of alpha-synuclein (αSyn) into cytoplasmic inclusions. This process is accompanied by elevated phosphorylation at serine 129 (S129). The accumulation of αSyn into aggregates and their propagation disrupts key proteostasis pathways, including the ubiquitin–proteasome system (UPS) or autophagy, contributing to cellular dysfunction and neuronal death. This study identified the proteasome activator Blm10 and its human ortholog PA200 as modulators of αSyn degradation and toxicity. The conserved Blm10/PA200 protein plays a key role in regulating proteasome activity and assembly. The αSyn expression increases Blm10 protein stability through autophagy inhibition, in a manner dependent on αSyn phosphorylation at S129 in yeast. Overexpression of *BLM10* or *PA200* reduces αSyn aggregation and enhances αSyn turnover via activation of the 20S proteasome in yeast and mammalian cells. Blm10 and PA200-capped 20S proteasomes efficiently degrade both monomeric as well as oligomeric αSyn *in vitro*. Notably, capped proteasomes retain proteolytic activities in presence of αSyn, indicating resistance to αSyn-induced inhibition, in contrast to 20S or 26S proteasomes. These results reveal a distinct proteasome subtype that bypasses UPS impairment and restores proteolytic capacity under proteotoxic stress. Our findings establish Blm10/PA200 as critical regulator of αSyn proteostasis and highlight its protective role in maintaining protein homeostasis and cell viability under conditions of αSyn toxicity.

## 1. Introduction

Proteasomal dysfunction is a hallmark of numerous neurodegenerative disorders, including Parkinson’s disease (PD). Proteasome activity declines with aging and can lead to the accumulation of misfolded α-synuclein (αSyn) protein aggregates. During disease progression, αSyn undergoes a conformational shift from a soluble, natively unfolded protein to β-sheet-rich oligomers and insoluble amyloid fibrils, which accumulate as Lewy bodies (LB), the pathological hallmark of PD (Spillantini et al., 1997; Winner et al., 2011; Mehra et al., 2021). This aggregation is related to a failure in protein clearance mechanisms, particularly the ubiquitin-proteasome system (UPS), which itself deteriorates with age (Hipp et al., 2019). Moreover, αSyn has been shown to impair both autophagy (Fellner et al., 2021) and proteasomal degradation (Lindersson et al., 2004; Thibaudeau et al., 2018), creating a pathological feedback loop that exacerbates proteotoxic stress. The precise molecular mechanisms connecting proteasome dysfunction to PD pathology remain poorly understood, although the inhibition of proteasomal activity has been linked to αSyn aggregation and neuronal loss. Elucidating these processes is essential for understanding disease progression and for identifying therapeutic targets within the UPS.

Soluble αSyn is primarily cleared by the UPS, whereas aggregated and oligomeric forms are preferentially degraded through autophagy (Webb et al., 2003; Petroi et al., 2012). Key regulatory determinants of αSyn turnover are its post-translational modifications, particularly phosphorylation at serine 129 (pS129), which is highly enriched in pathological aggregates (Anderson et al., 2006; Oueslati, 2016). This modification influences αSyn aggregation propensity, toxicity, and susceptibility to degradation, making it a focal point in PD pathophysiology (Shahpasandzadeh et al., 2014; Popova et al., 2015; Kleinknecht et al., 2016; Stefanis et al., 2019). The expression of αSyn disrupts proteasome homeostasis in cellular models by reducing the abundance of proteasomal subunits (McNaught et al., 2003; Popova et al., 2021a) and by impairing the assembly and activity of 26S proteasomes (Galka et al., 2024). αSyn degradation is mediated by different types of proteasomes, each with distinct roles. The 26S proteasome, which requires ATP and targets polyubiquitinated substrates, is primarily responsible for degrading soluble αSyn under normal conditions (Bi et al., 2021).

In contrast, the 20S core particle (CP) can directly degrade unstructured αSyn by ubiquitin- and ATP-independent mechanisms (Tofaris et al., 2001). This pathway is particularly relevant during oxidative stress when ubiquitination is compromised. Oxidative stress has been shown to occur in cells with αSyn pathology (Hsu et al., 2000). Additionally, hybrid proteasomes exist in cells, formed by the association of the 20S core with alternative regulatory particles that possess intermediate substrate specificities (Mayor et al., 2016). These variants could influence αSyn turnover during cellular stress or inflammation, however their specific contribution is not yet defined. Understanding the balance and regulation among these proteasomal subtypes is critical for elucidating αSyn homeostasis and its disruption in PD.

A recent high-throughput screen identified Blm10 as a protein stabilized in the presence of αSyn (Galka et al., 2024), suggesting a potential role in αSyn homeostasis. This study investigated how the proteasome interacting protein Blm10 modulates proteasome activity and facilitates αSyn degradation. Yeast Blm10 as well as its human ortholog PA200 bind the 20S proteasome and enhance the degradation of small, unfolded proteins (Schmidt et al., 2005; Dange et al., 2011; Weberruss et al., 2013). Expression levels of the *PSME4* gene which encodes PA200, as well as *BLM10,* decrease during aging, making them potentially interesting target for analysis in the context of PD (Chen et al., 2020). The functional contribution of Blm10 in counteracting αSyn toxicity was examined by using *Saccharomyces cerevisiae* as well-established reference cell system for PD (Outeiro and Lindquist, 2003; Petroi et al., 2012). Elevated Blm10 levels improve cell viability, accelerate αSyn turnover, and restore proteasome activity both *in vivo* and *in vitro*. Importantly, the effects observed in yeast proteasomes with Blm10 could be recapitulated by human 20S proteasomes capped with PA200. The findings demonstrate that Blm10/PA200-mediated 20S proteasome activation enhances αSyn degradation and mitigates proteasome dysfunction. These results reveal that targeting proteasome activation through subunit assembly or 20S gate opening mechanisms has a promising potential as therapeutic strategy to promote αSyn clearance.

## 2. Experimental Procedures

The yeast strains and plasmids used are listed in Table S1 and Table S2.

### 2.1 Yeast strains, transformations and culture conditions

Plasmids were constructed using GENEART seamless cloning and assembly kit (Invitrogen, USA) and verified by DNA sequencing. *S. cerevisiae* yeast strains were grown at 30 °C in YEPD (Yeast-Extract-Peptone-Dextrose) media or in synthetic complete (SC) dropout medium, lacking the respective amino acids for selection. The medium was supplemented with either 2% glucose, 2% raffinose, or 2% galactose. Expression of genes under *GAL1* promotor control was achieved by supplementing the SC medium with 2% galactose. Yeast transformations were performed using the standard lithium acetate procedure (Gietz et al., 1992) and plasmids isolated from DH5α *E. coli* bacteria. Human embryonic kidney 293 (HEK) cells were maintained as described previously (Popova et al., 2021b).

### 2.2 Fluorescence microscopy

Overnight cell cultures cultivated in raffinose-containing SC medium lacking amino acids for selection were diluted to OD_600_ = 0.3 and transferred to SC medium containing 2% galactose. After induction for 6 h, 300 µL cells were transferred into 8-well Ibidi Dishes (Ibidi, Germany) and imaged using a Zeiss Observer Z1 microscopy (Carl Zeiss, Germany) equipped with a CSU-X1 A1 confocal scanner unit (Yokokgawa, Japan) and a QuantEM:512SC digital camera (Photometrics, USA). Fluorescence intensities were measured using Slide Book 6.0 software package (Intelligent Imaging Innovations, USA).

### 2.3 Cycloheximide chase experiment

Genes coding for target proteins were expressed for 6 h before the cultures were diluted to equal cell density and supplemented with 50 µg/mL cycloheximide to stop translation. An equal volume of samples was collected at time points 0 h, 1 h, and 2 h. Cells were pelleted, frozen in liquid nitrogen and used for crude protein extraction.

### 2.4 Immunoblotting

Yeast cells were harvested by centrifugation and yeast protein extraction was performed by NaOH lysis and trichloroacetic acid (TCA) precipitation. The cell pellet was resuspended in 1 mL 0.25 M NaOH with 1.5% 2-mercaptoethanol for cell lysis. After 15 min incubation on ice, the proteins were precipitated with 150 µL 55% TCA. Protein pellets were resuspended in HU buffer (200 mM Tris-HCl pH 6.8; 8 M Urea; 5% SDS; 1 mM EDTA; 0.05% bromophenol blue) and denatured at 65°C for 10 minutes. Gel electrophoresis was performed in 9% or 12% SDS-acrylamide gels, depending on the size of the proteins of interest. Proteins were blotted onto a nitrocellulose membrane (GE Healthcare, USA) and western hybridization analyses were performed using standard procedures (Popova et al., 2021b). Antibodies used are listed in Table S3. Quantifications of pixel densities were obtained from TIFF files originating from digitized X-ray films (Cytiva, USA) and analysed with ImageJ software (NIH, USA).

### 2.5 Spotting assay

Spotting assays for analysis of yeast growth were performed using cells grown over night in selective SC medium supplemented with 2% raffinose. Cells were diluted to OD_600_ = 0.1 and serially diluted 10-fold. Cells were spotted in a volume of 10 µL onto selective SC plates containing glucose as a control or galactose to induce expression of genes under *GAL1* promotor. Plates were incubated at 30 °C and documented after three days.

### 2.6. Growth Analysis in liquid culture

Growth of yeast cells was analysed in liquid media after overnight growth in selective SC media containing 2% raffinose at 30°C in a clear 96-well plate in a Tecan Infinite M200 (Tecan, Switzerland) plate reader till all samples reached stationary phase. The next day 1 µL of each sample was transferred onto a new 96-well plate with selective SC medium containing raffinose. After 90 min 20% galactose was diluted in the cultures to a concentration of 2% in order to induce expression of genes under *GAL1* promotor. Cell growth at 30 °C was observed for 20 h by measuring the absorbance at 600 nm.

### 2.7. Peptidase activity measurement

Proteasome activities were measured through the degradation of the fluorogenic peptide SUC-LLVY-AMC (Enzo Life Science, USA) and the subsequent release of 7amino-4-methylcoumarin. Fluorescence was measured for 30 min at 37°C using Tecan Infinite M200 plate reader (excitation wavelength = 350 nm; emission wavelength = 440 nm). Crude protein extracts were harvested from transformed yeast cells that were grown in selective SC medium containing galactose for 16 h to induce expression of genes under *GAL1* promotor. Yeast cells were lysed by cryo-milling in buffer A (50 mM Tris-HCl pH 7.4; 100 mM NaCl; 10% Glycerol; 10 mM MgCl_2_; 4 mM ATP). Human epithelial kidney (HEK) cells were lysed by sonication in RIPA buffer (50 mM Tris-HCl pH 8.0; 150 mM NaCl; 0.1% SDS; 0.5% Na-deoxycholate; 1% NP-40) supplemented with protease inhibitor mix (PIM; Roche Diagnostics, Germany). Cell lysates were cleared by centrifugation at 4°C for 15 min at 15.000g. Concentration of protein extract was determined by Bradford assay and 100 µg crude protein extract were used for 26S activity assays using SUC1-buffer (20 mM Tris-HCl pH 7.4, 50 mM NaCl, 2 mM DTT, 5 mM MgCl_2_, 2mM ATP) supplemented with 100 µM SUC-LLVY-AMC. To assess the activity of 20S proteasomes in crude protein extracts, 0.025% SDS was added to the SUC1-buffer. For determination of the activity of *in vitro* reconstituted 20S+Blm10 or human 20S+PA200, purified 20S proteasomes were reconstituted with the respective proteasomal activator for 30 min at 30°C. To assemble the proteasomes, 250 nM proteasome and 500 nM Blm10 or PA200 were added to the reconstitution buffer (50 mM Tris-HCl pH 7.4; 5 mM MgCl_2_; 0.5 mM EDTA). 0.5 µg purified 20S proteasomes were used for activity assays. The peptidase activity of purified 20S proteasomes was measured using 100 µM SUC-LLVY-AMC in SUC2-buffer (50 mM Tris–HCl pH 7.4, 100 mM NaCl, 10 mM MgCl_2_, 2 mM DTT) as described above.

### 2.8. Protein purification

Proteasomes were purified using *RPN11-3xFLAG* and *PRE1-3xFLAG* strains for 26S and 20S proteasomes, respectively. Cells were grown in YPED media overnight, harvested, and cryo-milled in buffer A (for 26S) or buffer B (50 mM Tris-HCl pH 7.4; 500 mM NaCļ1 mM EDTA) for 20S proteasomes. The absence of ATP and increased NaCl concentration removes the regulatory particle from the core particle (Li et al., 2015). Human proteasomes were purchased commercially (Enzo Life Science, USA). Yeast cells transformed with plasmids coding for 3xFLAG-Blm10 were grown in selective SC media containing galactose to induce gene expression. Cells were harvested and cryo-milled in buffer B supplemented with protease inhibitor mix (PIM; Roche Diagnostics, Germany). Cell lysates were centrifuged for 30 min at 15 000 g at 4°C, and supernatant was used for FLAG affinity purification using the M2 affinity gel (Merck KGaA, Germany) according to the manufacturer protocol. For the purification of PA200, the protein was genetically fused to a 6xHis tag. The corresponding coding sequence was cloned into a yeast plasmid and expressed in yeast cells. Gene expression was induced with galactose overnight, after which cells were harvested. Cells were cryo-milled in native binding buffer (50 mM NaH_2_PO_4_ pH 8.0; 500 mM NaCl, 10 mM Imidazole) supplemented with protease inhibitor mix (PIM; Roche Diagnostics, Germany). Cell lysates were cleared by centrifugation at 4°C for 15 min at 15.000 g. Native protein purification of His-tagged PA200 was performed according to the Novex® Ni-NTA manufacturer protocol (Thermo Fisher Scientific, USA) using Ni-NTA agarose beads (Qiagen N.V., The Netherlands).

### 2.9. Expression and purification of human recombinant αSyn

Expression and purification of αSyn was performed as described previously (Miranda et al., 2013). *E. coli* strain BL21 (DE3) was transformed with pME4913 and the expression was induced in LB medium using 1 mM isopropyl-β-D-1-thiogalactopyranoside (IPTG). The overnight culture was inoculated at OD_600_ = 0.3, grown at 37°C, pelleted and flash-frozen at −80°C. Cells were lysed using by sonication (five steps, 30 s per step, 1 min cool down in-between steps) in *E.coli* lysis buffer (750 mM NaCl, 10 mM Tris pH 8.0, 1 mM EDTA, 1 mM PIM). Afterwards the lysate was heated to 95 °C for 15 min and cleared by centrifugation at 4 °C for 10 min at 15.000 g. Supernatant was dialyzed overnight at 4 °C against the dialysis buffer (50 mM NaCl, 10 mM Tris pH 7.6, 1 mM EDTA). Following this, αSyn was purified using HiTrap Q FF 1 mL anion exchange columns (GE Healthcare, USA) using a NaCl gradient from 0-600 mM in 25 mM TrisHCl pH 7.7. Fractions were collected and analysed using SDS-PAGE and fractions containing αSyn were further purified using a Superdex 75 26/600 prep grade 120 mL size-exclusion chromatography column. Elution was performed in SEC buffer (100 mM NaCl, 25 mM HEPES, 1 mM DTT equilibrated to pH 8.0). The resulting fractions were analysed for αSyn purity and aggregation state using SDS-PAGE and immunoblot analysis, protein concentration was determined by Bradford assay, and proteins were stored at 181 cl:153780°C.

### 2.10. Native PAGE and in-gel activity

Native PAGE with following in-gel activity measurement was performed with reconstituted 20S proteasomes. Gel electrophoresis was performed using 4% acrylamide gel complemented with 5 mM MgCl_2_ and 2.5% sucrose and 3% stacking gel. Electrophoresis was run for 3 h on ice at 100 V using filter sterilized native running buffer (90mM Tris-Borate pH 8.3; 5 mM MgCl_2_). In-gel peptidase activity was measured after incubating the gel for 10 min at 37°C using SUC2-buffer supplemented with 100 µM SUC-LLVY-AMC. The resulting fluorescent activity was captioned using Fusion FX6 Edge Imaging System (Vilber, France) using the corresponding software. Signal intensity in the resulting TIFF files was measured using Image J (NIH, MD, USA).

### 2.11. Negative stain Transmission Electron Microscopy

Purified 20S proteasomes where incubated with purified Blm10 at indicated concentrations for 30 min at 30 °C. For negative stain analysis, samples were loaded on a continuous carbon film. The copper grid (CF200, Electron Microscopy Sciences, PA, USA) was glow-discharged using a plasma cleaner (HARRICK PLASMA, USA). The carbon film was attached to the grid and stained in 2% Uranyl-acetate solution (Science Services) for 1 min. Afterwards the sample was dried and imaged using Tecnai G2 spirit (Thermo Fisher Scientific, USA).

### 2.12. *In vitro* degradation of αSyn

Purified αSyn protein was incubated with 20S proteasomes or 20S+Blm10 or 20S+PA200 complexes, reconstituted as described above to analyse the proteasomal degradation of αSyn. For each time point 0.5 µg of αSyn was added to 2.4 µg of proteasomes. Degradation of αSyn occurred at 30 °C over a two-hour time frame and samples were taken at the indicated time points and denatured at 95 °C for 10 min in Laemmli buffer (62.5 mM Tris-HCl pH 6.8; 2% SDS; 2.5% β-mercaptoethanol, 5% glycerol; 0.005% bromophenol blue). Degradation of αSyn was monitored using SDS-PAGE with subsequent immune-hybridization analysis.

### 2.13. Human cell culture and transfection

Human neuroglioma H4 cells were cultured at 37 °C and 5% CO₂ in Opti-MEM Reduced Serum Medium (Life Technologies-Ginco, Carlsbad, CA), supplemented with 10% fetal bovine serum (FBS) (PAA, Cölbe, Germany) and 1% Penicillin-Streptomycin. Twenty-four hours prior to transfection, cells were seeded into 12-well plates (Costar, Corning, New York). Cells were transfected with FuGENE® (Promega, WI, USA) according to the manufacturer’s protocol. Plasmids encoding SynT and synphilin-1 were co-transfected at a 1:1 ratio, or together with PA200 at a 1:1:1 ratio. 48 h post-transfection, cells were fixed for immunostaining.

### 2.14. Immunocytochemistry

Cells were fixed with 4% paraformaldehyde in DPBS and washed three times. Permeabilization was performed for 10 min with 0.1% Triton X-100 in DPBS, followed by blocking for 1 h at room temperature with 1.5% normal goat serum (S1000, Vector). Cells were incubated overnight at 4 °C with primary antibodies: mouse anti-α-synuclein (Syn1, 1:1000; BD Transduction Laboratories) and rabbit anti-FLAG (1:1000; F-7425, Sigma-Aldrich). After PBS washes, cells were incubated for 1 h at room temperature with Alexa Fluor 488 donkey anti-mouse IgG and Alexa Fluor 555 goat anti-rabbit IgG (both 1:1000; Invitrogen). Nuclei were stained with DAPI (Carl Roth, Germany) for 5 min. Coverslips were mounted using Fluoromount-G (Invitrogen) and stored at room temperature until imaging.

### 2.15. Statistical analysis

Quantitative data were analysed using GraphPad Prism 6 software (San Diego, USA) and presented as means ± SEM of at least three individual replicates. Statistical significance was assessed as described in the figure legend, with p < 0.05 considered significant.

## 3. Results

### 3.1. αSyn-induced autophagy impairment enhances the stability of the 20S proteasome activator Blm10

A proteome-wide screen using tandem fluorescent protein timer (tFT) fusions was previously conducted to explore changes in protein stabilities upon expression of αSyn or the phosphorylation-deficient S129A mutant in yeast (Galka et al., 2024). tFT construct consists of a tandem fusion of mCherry and superfolder GFP (sfGFP), fluorescent proteins with distinct kinetics of fluorophore maturation. This enables direct visualization of protein stability through colour changes over time. Expression of αSyn led to significant disruption of protein homeostasis and revealed a novel connection between αSyn and 26S proteasome assembly. The stability of the proteasomal activator Blm10-tFT increased by expression of αSyn or the phosphorylation-deficient S129A variant increased. Blm10 emerged as top candidate among the stabilized proteins. Live cell fluorescence microscopy of *BLM10*-tFT yeast strain showed stronger Blm10 stabilization upon αSyn expression compared to S129A (Figure 1a, b). Expression of the S129D variant protein that mimics constant phosphorylation at S129 further confirmed that αSyn phosphorylated at S129 affects Blm10 stability significantly more than the non-phosphorylatable S129A version. Similarly, hyperphosphorylation of αSyn through overexpression of the human kinase GRK5 produced a comparable effect (Supplemental Figure S1). This kinase is known to phosphorylate αSyn at S129 in humans as well as in yeast (Shahpasandzadeh et al., 2014). Blm10, like its mammalian ortholog PA200, contains a conserved C-terminal HbYX motif that promotes 20S proteasome gate opening and enhances proteolytic activity (Chuah et al., 2023). Both Blm10 and PA200 promote ubiquitin-independent degradation of unfolded, aggregation prone proteins such as tau and huntingtin *in vitro* (Aladdin et al., 2020), and have been shown to mitigate proteasomal inhibition caused by these proteins (Chuah et al., 2023). Next, the question was addressed, whether the effect of αSyn or S129A on Blm10 stability is mediated by direct physical interaction between the proteins using two different approaches. Yeast-two-hybrid (Y2H) (Figure S2) and Bimolecular fluorescence complementation (BiFC) experiments (Figure S2b, c) revealed that there is no direct interaction between the proteins. These findings suggest that the impact of αSyn on Blm10 stability is likely mediated by indirect effects rather than direct physical interaction.

**Figure 1.**
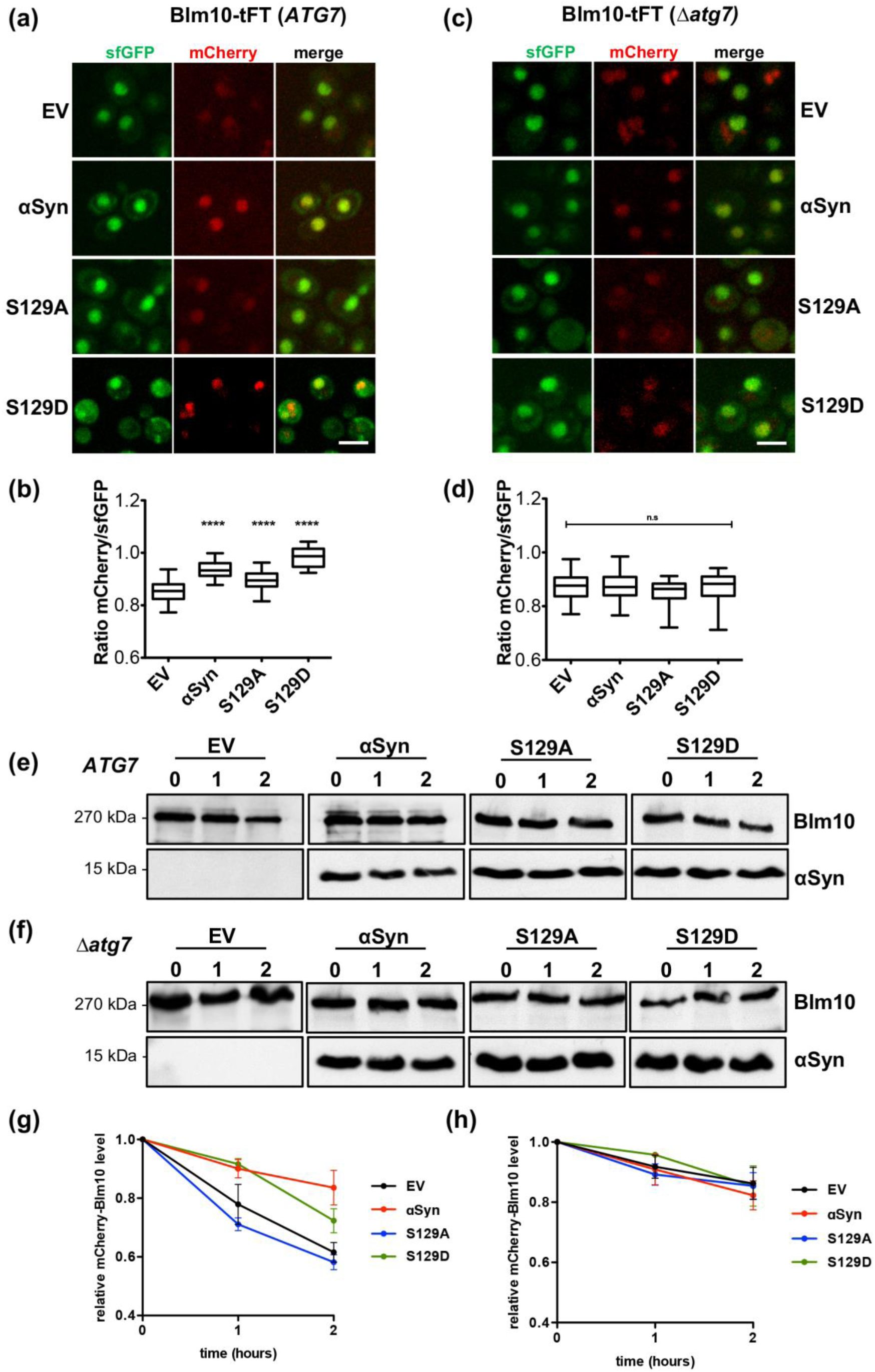
αSyn expression increases Blm10 stability via autophagic inhibition. **(a)** Tandem Fluorescent Timer (tFT) analysis of Blm10 stability by fluorescence microscopy. Fast-maturing sfGFP and slow-maturing mCherry allow estimation of protein stability via mCherry/sfGFP fluorescence ratios. αSyn or variants expressed for 6 h increase Blm10 stability versus empty vector (EV). Scale bar = 5 µm. **(b)** Quantification of single-cell mCherry/sfGFP ratios; one-way ANOVA with Dunnett’s post hoc (****p < 0.0001; n = 50) versus EV. **(c, d)** tFT analysis in Δ*atg7* cells shows no significant Blm10 stability differences with αSyn or variants. Scale bar = 5 µm. **(e, f)** Immunoblots of mCherry-Blm10 degradation after cycloheximide (CHX) chase in *ATG7* and Δ*atg7* cells. **(g, h)** Quantification of Blm10 levels at indicated times post-CHX in *ATG7* (g) and Δ*atg7* (h) cells.

Under conditions of physiological *BLM10* expression, Blm10 is degraded predominantly by autophagy (Burris et al., 2021). αSyn negatively influences both the selective and nonselective autophagy pathways (Sahoo et al., 2022). Therefore, it was assessed whether Blm10 stabilization by αSyn is due to reduction of its degradation through the autophagy pathway. The impact of αSyn variants on autophagy was tracked using the autophagy marker GFP-Atg8 which associates with autophagosomal membranes and enables visualization of autophagosomes. Expression of αSyn variants coincided with accumulation of autophagosomes in the cytoplasm, indicating inability of autophagosomes to fuse with the vacuole of yeast cells (Figure S3a, b). Additionally, the impact of αSyn and S129 phosphorylation on autophagy flux was assessed. Autophagy was induced by nitrogen starvation and the cleavage of GFP from GFP-Atg8 upon delivery to the vacuole was monitored (Figure S3c, d). The results revealed that expression of αSyn inhibits autophagy in dependence of S129 phosphorylation. The effect of αSyn on Blm10-tFT stability was further investigated in the Δ*atg7* deletion strain. *ATG7* encodes a protein that is required for autophagy as it activates the ubiquitin-like proteins Atg8 and Atg12, which are indispensable for autophagosome formation. Inability of cells to perform autophagy was verified in the presence of αSyn variants and in control cells (Figure S3e). No difference in Blm10-tFT stability was observed between the control and αSyn or its variants when autophagy is inhibited (Figure 1c, d), indicating that autophagic degradation is the major regulatory pathway of Blm10 stability affected by αSyn expression.

Cycloheximide chase experiments using N-terminally tagged mCherry-Blm10 further confirmed that the change in Blm10 stability is caused by reduced autophagic degradation (Figure 1e-h). Cells harvested at indicated time points were used for immune-hybridization and the degradation of Blm10 was followed over time. In cells with intact *ATG7*, expression of αSyn led to increased stability of mCherry-Blm10 (Figure 1e, g). The inhibition of Blm10 degradation was higher in cells with phosphorylatable S129, corroborating the results from Blm10-tFT experiments. In the Δ*atg7* strain no significant differences in Blm10 degradation were observed (Figure 1f, h). In summary, the presence of αSyn impairs the degradation of Blm10 through inhibition of autophagy. Since the inhibition of autophagy is dependent on the phosphorylation of S129, the increased stability of Blm10 is also linked to the phosphorylation of this residue.

### 3.2. Elevated *BLM10*/*PSME4* expression levels reduce αSyn aggregation in yeast and mammalian cells and improve yeast cell growth

Given the observed increase in Blm10 stability in the presence of αSyn, it was investigated whether elevated levels of Blm10 modulate αSyn-associated toxicity and aggregation and how pS129 contributes to this process. αSyn, S129A were co-expressed with mCherry-*BLM10*, testing different levels of *BLM10* by expression from low-copy (*CEN*) or high-copy (2µ) plasmids. Growth tests were performed to determine αSyn or S129A-associated cytotoxicity. High-level overexpression of *BLM10* improved the growth of yeast cells expressing either wild-type αSyn or the S129A variant (Figure 2). Low-level overexpression from a *CEN* plasmid conferred minimal or no protective effect compared to endogenous *BLM10* expression. Deletion of *BLM10* did not significantly affect the growth of αSyn expressing cells, suggesting that Blm10 may partially rescue αSyn toxicity when expressed at supraphysiological levels. Fluorescence microscopy was used to determine whether Blm10 influences cytotoxicity associated αSyn-GFP aggregation. Cells expressing αSyn-GFP exhibited prominent cytosolic aggregates (Figure 2b, c). In contrast, Blm10 overexpressing cells showed a significant reduction in αSyn-GFP aggregation even at moderate overexpression levels.

**Figure 2.**
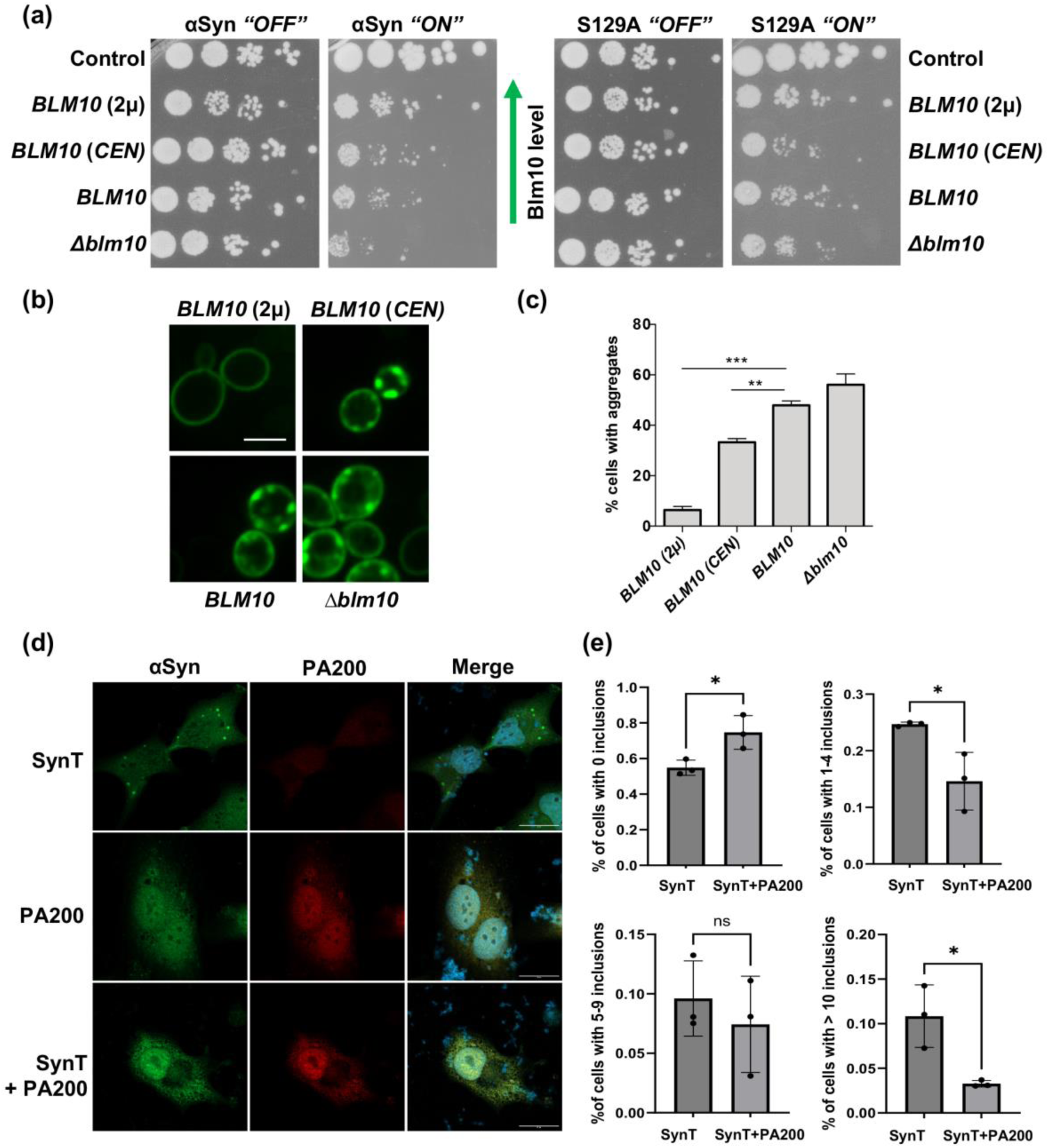
Increased Blm10/PA200 levels reduces αSyn aggregation in yeast and mammalian cells. **(a)** Yeast growth assays with increasing levels of Blm10. *BLM10* is expressed from low-copy (*CEN*) or high-copy (2µ) plasmids with *GAL1*-driven wild-type αSyn or S129A for 3 days. **(b)** Fluorescence microscopy of yeast αSyn-GFP after 6 h; foci form at the plasma membrane. **(c)** Quantification of αSyn-GFP foci; one-way ANOVA with Dunnett’s post hoc versus endogenous *BLM10* (***p < 0.001; **p < 0.01; n = 3). **(d)** Fluorescence microscopy of human H4 cells transfected with synphilin-1 and synT ± PA200, immunostained for αSyn and PA200. Scale bar = 30 µm. **(e)** Quantification of cytoplasmic inclusions per cell (0, 1–4, 5–9, >10); mean ± SD; Student’s t-test (*p < 0.05; n = 3).

Next, we assessed whether elevated levels of PA200 inhibit αSyn aggregation in human neuroglioma (H4) cells. αSyn aggregation can be induced by co-expression of a C-terminally modified form of αSyn (SynT) and synphilin-1, an αSyn-interacting protein that facilitates the formation of intracellular αSyn inclusions (Lázaro et al., 2014). Therefore, H4 cells were co-transfected with plasmids encoding SynT, synphilin-1, and PA200. Similar to the finding from yeast cells, overexpression of *PSME4* reduced the percentage of cells with αSyn inclusions as well as the number of inclusions per cell (Figure 2d, e). These results corroborate that elevated levels of Blm10/PA200 rescued αSyn-associated growth retardation in yeast and reduced αSyn aggregation in yeast and in mammalian cells, respectively. The findings highlight the protective role of Blm10/PA200 in mitigating αSyn aggregation and toxicity. This corroborates that enhancing Blm10/PA200 function can provide a promising potential therapeutic strategy to counteract αSyn-induced cellular dysfunction.

### 3.3 *BLM10* overexpression promotes αSyn clearance by reducing steady-state levels and enhancing its turnover

Previous studies have demonstrated that αSyn exerts a dose-dependent cytotoxic effect in yeast, with higher cytosolic levels correlating with increased aggregate formation and enhanced toxicity (Petroi et al., 2012). Independently, Blm10–CP assemblies have been shown to facilitate degradation of small, misfolded proteins such as Tau and Huntingtin (Dange et al., 2011; Aladdin et al., 2020). Steady-state αSyn protein levels were analysed using immuno-hybridization to assess whether the protective effect of Blm10 in αSyn-expressing cells is linked to a reduction of αSyn burden (Figure 3a, b). Overexpression of *BLM10* significantly reduced cellular αSyn levels, even at lower expression levels. A similar effect was observed in cells expressing the phosphorylation-deficient S129A variant of αSyn, although the effect was notably attenuated (Figure 3c, d).

**Figure 3.**
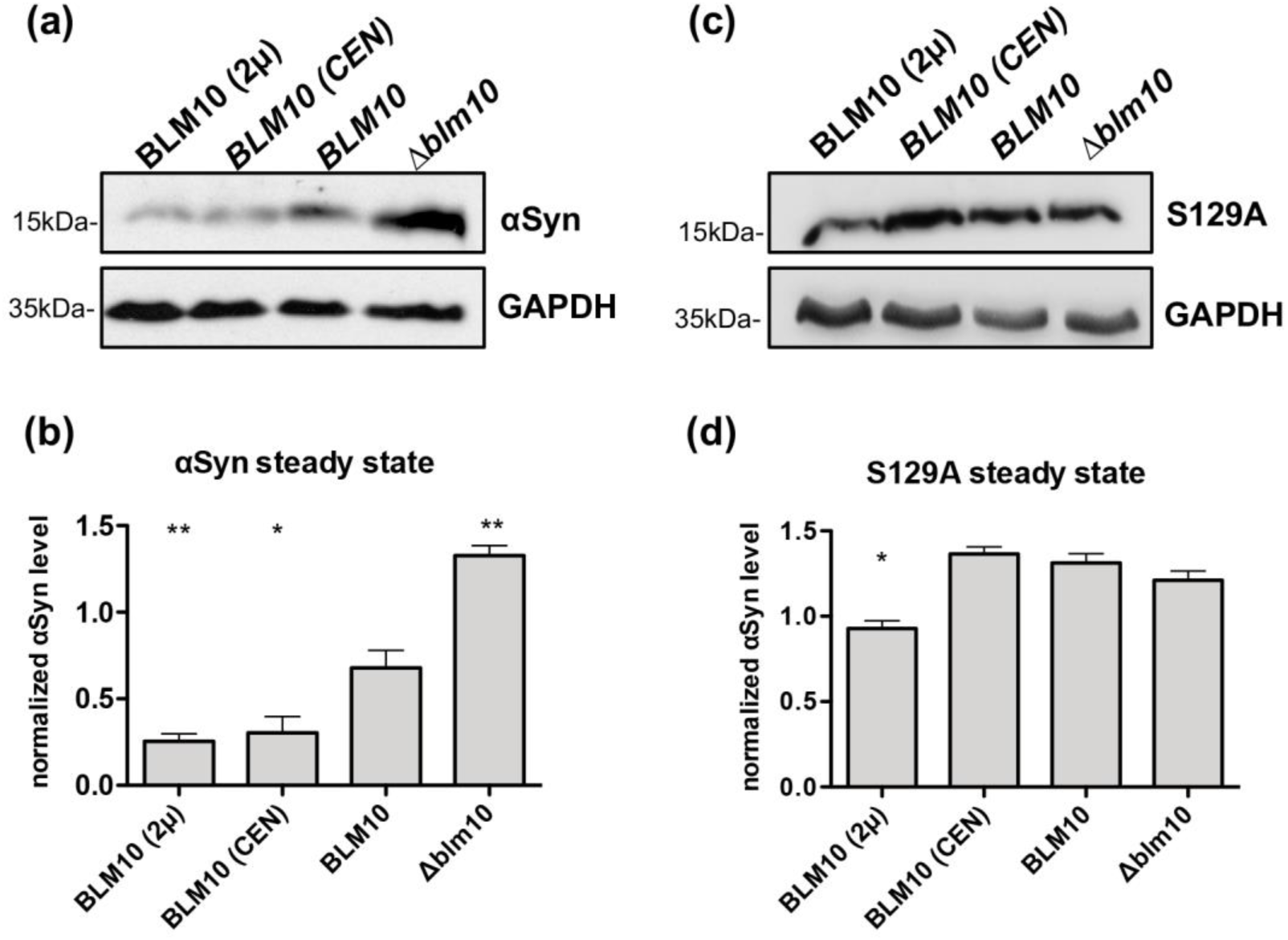
Elevated Blm10 levels reduce αSyn steady-state levels. **(a, c)** Immunoblots of αSyn or S129A after 6 h *GAL1*-driven expression with varying *BLM10* expression levels. **(b, d)** Quantification of αSyn or S129A levels normalized to GAPDH; Student’s t-test relative to endogenous *BLM10* (**p < 0.01; *p < 0.05; n = 3).

Promoter shut-off assays were performed to determine whether the reduction in αSyn levels was due to increased protein turnover. Expression of αSyn-encoding *SCNA* gene was driven by the yeast *GAL1* promoter, which is effectively repressed upon glucose addition. Following glucose-induced transcriptional repression, αSyn degradation was monitored by immunohybridization at indicated time points (Figure 4). Cells overexpressing *BLM10* exhibited markedly accelerated αSyn degradation compared to those expressing it at endogenous levels (Figure 4b), which correlated with enhanced cell growth during the shut-off period (Figure 4c). This suggests that *BLM10* overexpression promotes αSyn clearance, thus mitigating cytotoxicity. The effect of *BLM10* overexpression was diminished in cells expressing the non-phosphorylatable S129A variant (Figure 4d). Specifically, clearance rates did not significantly differ across *BLM10* expression levels, and cell growth remained largely unaffected by *BLM10* expression levels after promoter shut-off (Figure 4e, f). These findings suggest that phosphorylation at serine 129 influences αSyn degradation by Blm10 containing proteasome assemblies. Collectively, these results indicate that increased expression of *BLM10* reduces αSyn levels of by promoting its degradation, potentially through enhanced assembly of Blm10-CP proteasomes, which emerge as prominent candidate for mediating proteasomal αSyn degradation.

**Figure 4.**
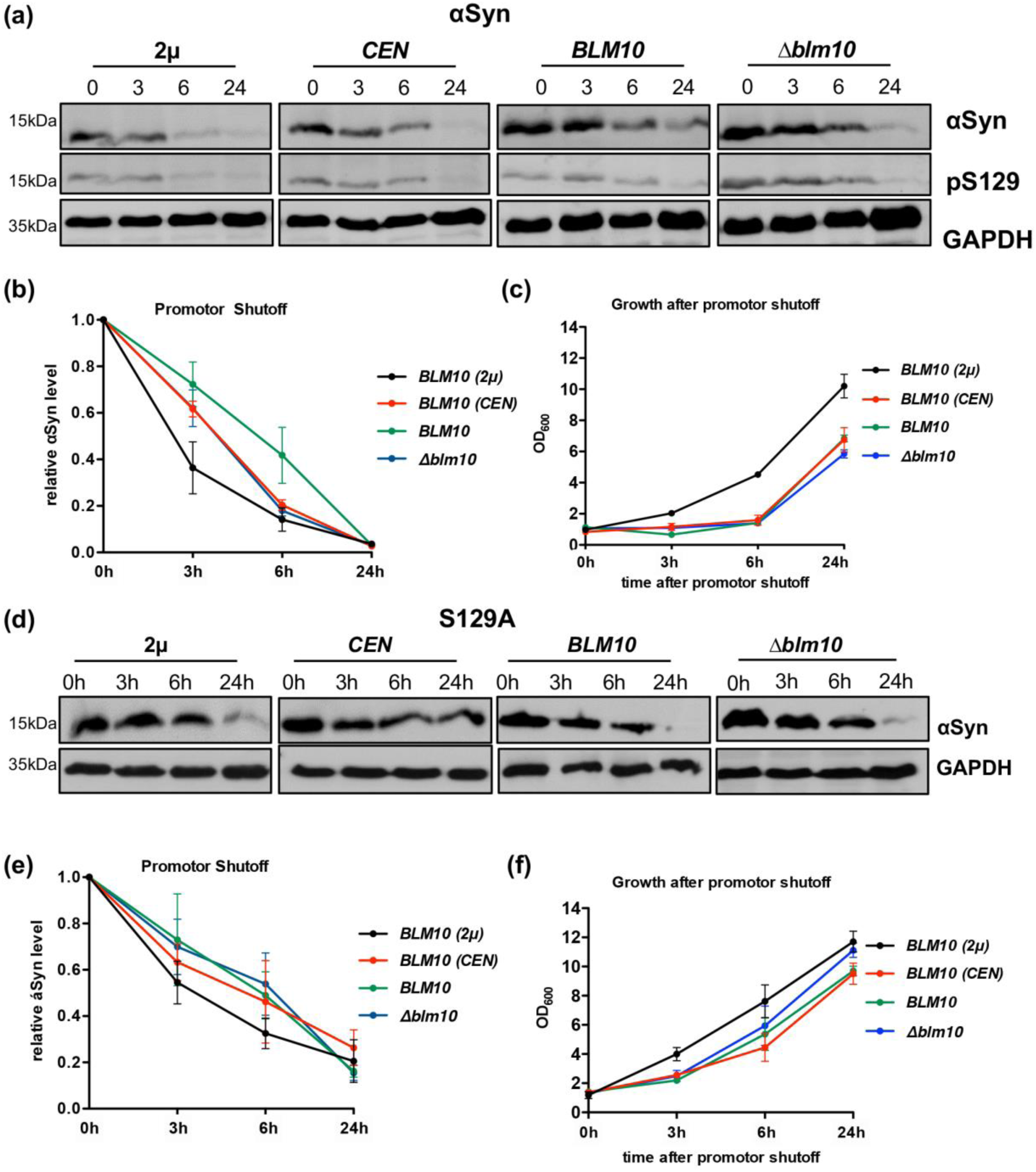
*BLM10* overexpression enhances αSyn turnover. **(a, d)** Immunoblots of αSyn (a) or S129A (d) protein levels after promoter shut-off at different *BLM10* expression levels. αSyn expression was induced for 6 h, then halted by addition of glucose. Samples were collected at indicated times; low-copy (*CEN*) and high-copy (2µ) *BLM10*. **(b, e)** Quantification of αSyn (b) or S129A (e) relative to 0 h. **(c, f)** Yeast growth in liquid culture post promoter shut-off.

### 3.4 Overexpression of *BLM10*/*PSME4* rescues αSyn induced proteasome inhibition *in vivo*

The interplay of αSyn expression and elevated Blm10 levels on the 26S or 20S proteasome assembly and resulting activities were examined. Crude protein extracts from yeast strains expressing different levels of *BLM10* were compared. The chymotrypsin-like proteasome activity was measured by monitoring the degradation of the fluorogenic peptide SUC-LLVY-AMC in the presence or absence of ATP and Mg²⁺. This approach allowed to distinguish between ATP-dependent 26S and ATP-independent 20S activities. The contribution of Blm10 was assessed by measuring proteasome activity under ATP-free conditions, where Blm10 specifically activates the latent 20S core particle by promoting gate opening. Assessment of 26S proteasome activity revealed a marked reduction in proteolytic function upon αSyn expression (Figure 5a). This inhibitory effect was less pronounced in cells expressing the S129A variant, indicating that phosphorylation at serine 129 contributes to αSyn-mediated 26S inhibition. Overexpression of *BLM10* partially restored 26S proteasome activity in cells expressing αSyn. In contrast, *BLM10* expression had minimal impact on 26S activity in the presence of S129A, further indicating that the Blm10 rescue effect is dependent on the specific αSyn post-translational modification. αSyn induced inhibition of 20S activity was less pronounced (Figure 5b). Importantly, overexpression of *BLM10*, even at moderate levels, robustly restored 20S activity in cells expressing αSyn. This effect was not observed in S129A-expressing cells, where 20S activity remained largely unaffected irrespective of *BLM10* expression levels. Taken together, these results suggest that αSyn impairs proteasomal activity, particularly that of the 26S complex, in a pS129-dependent manner. Whereas overexpression of *BLM10/PSME4* moderately alleviates 26S inhibition, it almost completely rescues the proteolytic impairment of the 20S proteasome.

**Figure 5.**
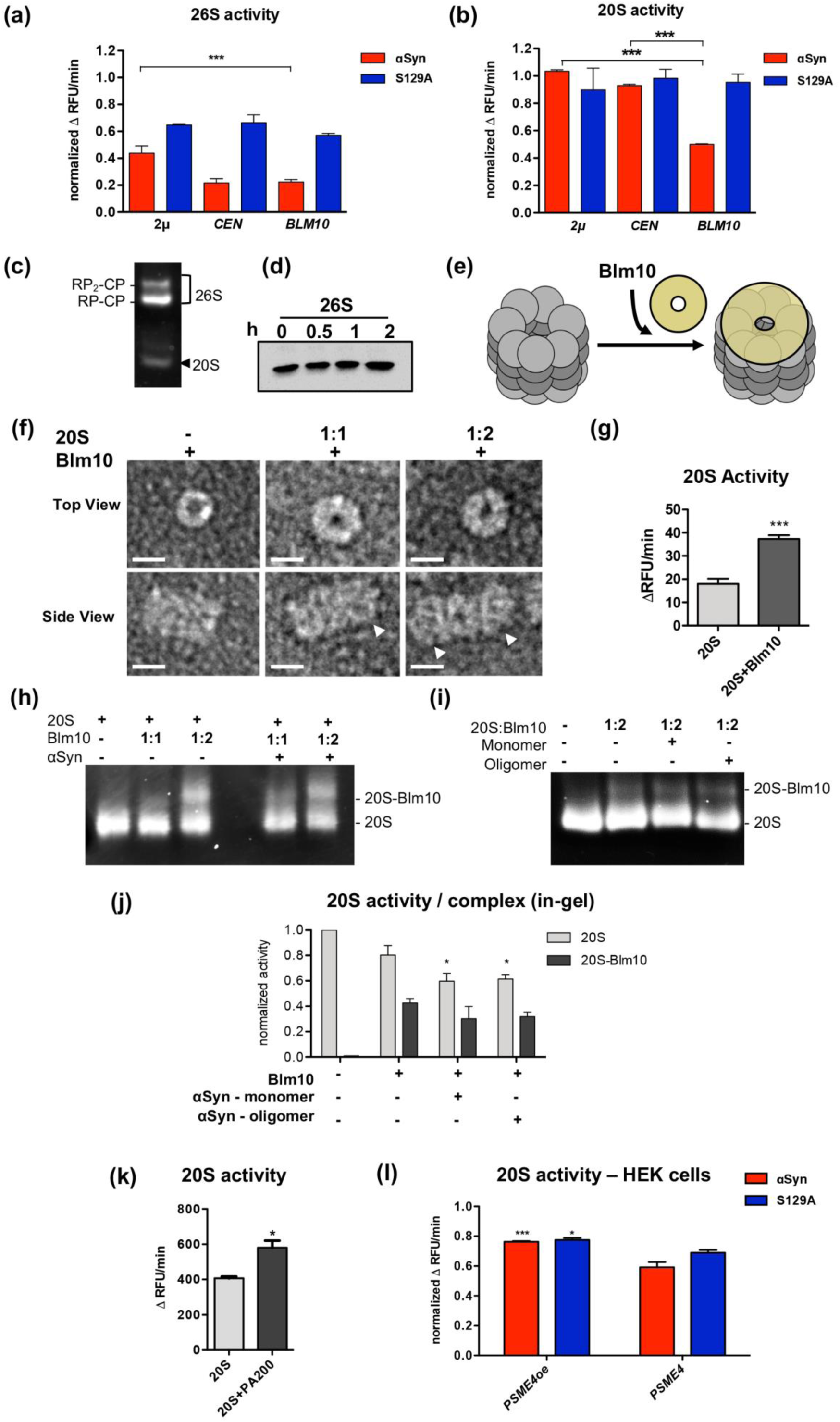
Blm10 rescues αSyn-induced proteasome inhibition and promotes proteasomal degradation. **(a, b)** 26S and 20S proteasome activity measured by SUC-LLVY-AMC assay in crude extracts from cells expressing αSyn or S129A with varying *BLM10* expression levels. Activity is presented as average change in relative fluorescence units (RFU) per minute and normalized to empty vector (EV) control for each *BLM10* condition. One-way ANOVA with Dunnett’s post hoc (***p < 0.001; n = 3). **(c)** In-gel proteasome activity following native gel electrophoresis of 3×FLAG-purified 26S proteasomes with ATP and 0.05% SDS, revealing single/double-capped 26S and uncapped 20S. RP – regulatory particle. **(d)** Immunoblot of *in vitro* αSyn degradation by 26S proteasomes at indicated time-points. **(e)** Schematic of 20S reconstitution with Blm10. **(f)** Negative-stain transmission electron microscopy of 20S ± Blm10 at varying stoichiometric ratios. White arrows indicate Blm10 caps on 20S particles. Scale bar = 10 nm. **(g)** SUC-LLVY-AMC assay of purified 20S ± Blm10 (molar ratio 1:2); Student’s t-test (**p < 0.01; n = 3). **(h)** In-gel SUC-LLVY-AMC activity assay following native gel electrophoresis, showing 20S proteasomes in complex with Blm10 ± 500 nM αSyn monomers. **(i)** In-gel SUC-LLVY-AMC activity of 20S + Blm10 at 1:2 ratio ± 500 nM αSyn monomers or oligomers. **(j)** Quantification of (i) normalized to 20S-only control. One-way ANOVA with Dunnett’s post hoc test (*p < 0.05; n = 3). **(k)** SUC-LLVY-AMC assay of purified 20S ± PA200 (1:2); Student’s t-test (*p < 0.05; n = 3). **(l)** 20S activity in crude extracts from HEK cells ± *PSME4* overexpression (oe); one-way ANOVA with Dunnett’s post hoc test (*p < 0.05; ***p < 0.001; n = 3).

*In vitro* experiments were performed to compare the ability of 20S, 26S, and hybrid proteasomes to degrade αSyn. Intact 26S proteasomes were isolated from yeast cells by affinity purification using *RPN11-3xFLAG* strain. The activities of the 30S double-capped (RP_2_-CP), 26S single-capped (RP-CP) or 20S proteasomes were visualized using Native PAGE followed by in-gel activity assay (Figure 5c). The efficacy of 26S proteasomes to degrade recombinant αSyn *in vitro* was analysed (Figure 5d). The degradation assay revealed that the 26S proteasome is unable to break down αSyn in the observed time frame in absence of additional co-factors, in accordance with recent findings (Maestro-López et al., 2025). To analyse whether Blm10-capped 20S core particle is able to degrade unfolded αSyn, we reconstituted the core particle (CP) with Blm10 (Figure 5e). 20S and Blm10 protein were isolated from yeast by affinity purification using *PRE1-3xFLAG* strain and *3xFLAG-BLM10* expressing cells, respectively. 20S-Blm10 proteasomes were reconstituted *in vitro* using 1:1 and 1:2 molar ratios of 20S to Blm10. The complex formation was analysed with negative-stain transmission electron microscopy (TEM) (Figure 5f). Complex formation at molar ratio 1:1 resulted in predominantly single-capped 20S-Blm10 complexes and uncapped 20S particles, whereas at molar ratio 1:2 most of the complexes were double-capped (Figure S4). Measurements of the enzymatic activity of the reconstituted complex validated that Blm10-capped proteasomes have higher activities compared to uncapped 20S proteasomes (Figure 5g).

Native PAGE and in-gel chymotrypsin activity assays also demonstrated successful reconstitution and activities of the 20S-Blm10 proteasomes (Figure 5h). Presence of recombinant αSyn did not inhibit the activity of 20S-Blm10 complex. It was further analysed whether αSyn oligomers might affect the assembly or activity of 20S-Blm10 complexes. αSyn oligomers were purified using size exclusion chromatography (Figure S5). The oligomeric fraction was analysed with Coomassie staining and dot-blot analysis using the oligomer specific A11 antibody (Figure S5b, c). The reconstitution mix was incubated with αSyn monomers or oligomers and subsequently loaded onto a native PAGE gel. SUC-LLVY-AMC hydrolysis shows that the activity of 20S-Blm10 complex remains unchanged in the presence of αSyn monomers or oligomers (Figure 5i, j). However, a reduction in 20S activity was visible in the αSyn containing samples.

The impact of αSyn monomers and oligomers on 20S proteasomes was further analysed using SUC-LLVY-AMC hydrolysis assay *in vitro*, revealing a dose dependent inhibition of 20S proteasomes by αSyn monomers or oligomers (Figure S6a, b). The inhibition of activity could be partially rescued when 20S proteasomes were reconstituted with Blm10 (Figure S6c-e). Similarly, human 20S proteasomes were reconstituted with PA200, purified from yeast (Figure 5k). Reconstituted human proteasomes exhibited significantly higher activity compared to uncapped 20S proteasomes. PA200 reconstitution partially alleviated the inhibition induced by αSyn monomers or oligomers (Figure S6f), although the effect was less pronounced than with yeast 20S proteasomes reconstituted with Blm10. Similar results were obtained when analysing 20S activity in HEK cell crude protein extracts (Figure 5l). Expression of αSyn or the phosphodeficient S129A variant significantly reduces 20S activity, which can be partially rescued by overexpression of *PSME4.* These results reveal that αSyn affects the activity of the 20S proteasome in its uncapped form, whereas Blm10-capped 20S complexes retain their proteolytic function even in the presence of αSyn monomers and oligomers, highlighting Blm10/PA200 protective role against αSyn-induced proteasomal impairment.

### 3.5. Blm10/PA200 capping facilitates αSyn clearance

Purified recombinant αSyn was incubated with either 20S proteasomes or Blm10 reconstituted 20S proteasomes (20S-Blm10 complexes) to determine whether the previously observed rescue effect of Blm10 on 20S activity is due to enhanced αSyn degradation. The efficiency of αSyn degradation was analysed by immuno-hybridization (Figure 6a), and quantified over time (Figure 6b). The 20S proteasomes displayed only limited ability to degrade αSyn, although their efficacy exceeded that of the 26S proteasomes (Figure 5d). In contrast, Blm10-capped 20S complexes exhibited markedly enhanced proteolytic activity, removing most of αSyn within one hour. These findings demonstrate that Blm10 significantly enhances the proteolytic activity of the 20S proteasome to degrade monomeric αSyn.

**Figure 6.**
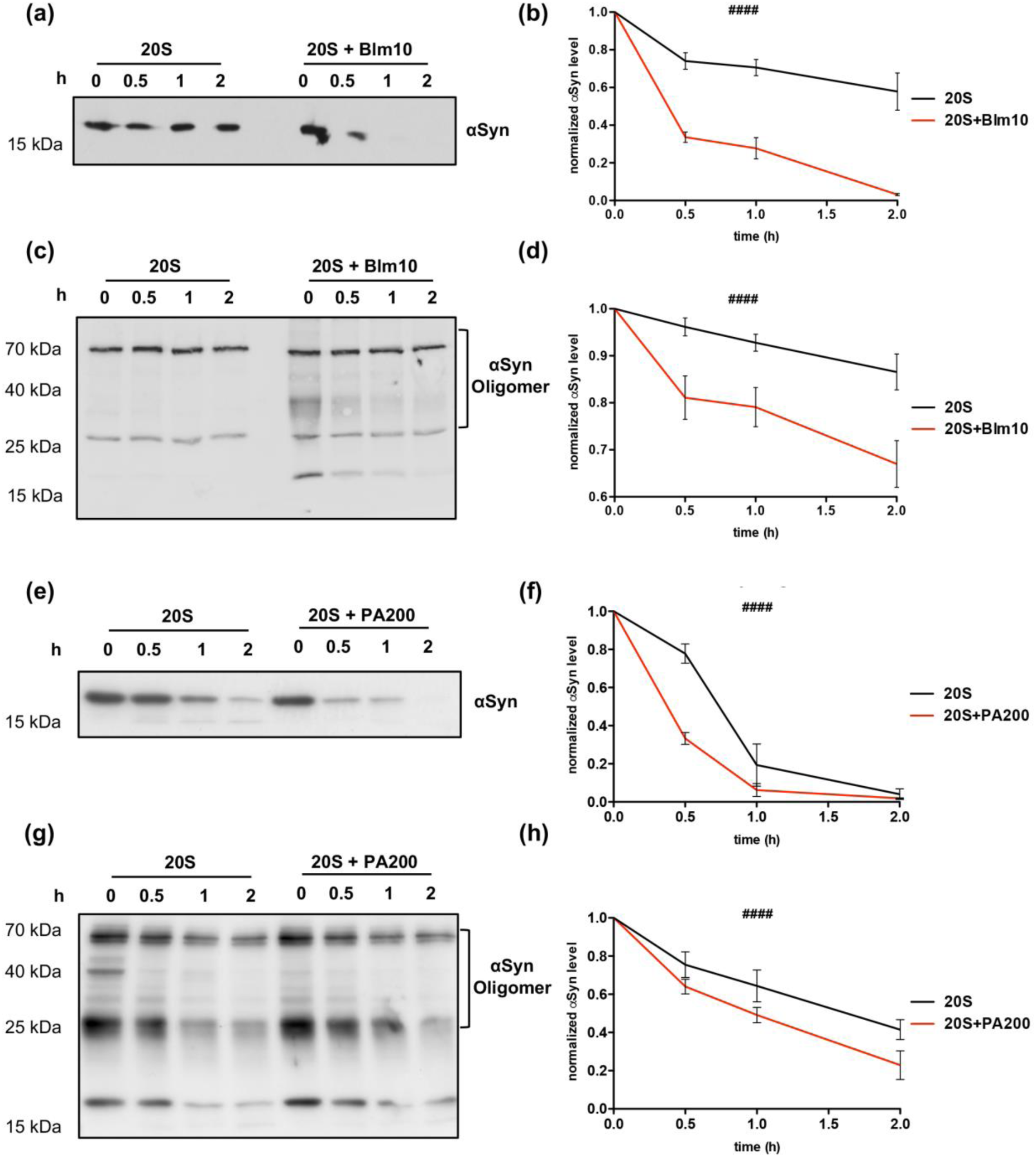
20S-Blm10 proteasome efficiently degrades αSyn in vitro. **(a, c)** Immunoblots of αSyn monomer (a) or oligomer (c) degradation over time by yeast 20S ± Blm10. Reactions contained 2.5 μg purified 20S, 2× molar excess of Blm10, and 0.5 μg αSyn at the reaction start. **(b, d)** Quantification of αSyn monomer (b) or oligomer (d) degradation normalized to time 0. **(e, g)** Immunoblots of αSyn monomer (e) or oligomer (g) degradation by human 20S ± PA200. Reactions contained 1 μg commercially purified 20S, 2× molar excess of PA200, and 0.5 μg αSyn at the reaction start. **(f, h)** Quantification of αSyn monomer (f) or oligomer (h) degradation normalized to time 0; two-way ANOVA (#### p < 0.0001; n = 3).

Given that A11-positive oligomers have been identified as major contributors to αSyn mediated cytotoxicity and proteasome inhibition (Winner et al., 2011; Thibaudeau et al., 2018), we assessed the ability of Blm10-capped proteasomes to degrade these oligomers (Figure 6c, d). Blm10-capped 20S proteasomes moderately degraded αSyn oligomers *in vitro* into smaller assemblies. Degradation products of approximately 40 kDa and 25 kDa appeared at the earliest time point and were partially degraded over the two-hour incubation. In contrast, uncapped 20S proteasomes neither reduced oligomer levels nor generated detectable degradation products.

To validate the findings with Blm10-capped 20S proteasomes, PA200 was reconstituted with human 20S proteasomes, and αSyn degradation was assessed similarly. PA200-capped 20S proteasomes degraded αSyn monomers significantly more efficiently *in vitro*, although the difference between capped and uncapped proteasomes was less pronounced than in yeast proteasomes (Figure 6e, f). Moreover, the degradation of αSyn oligomers was markedly enhanced in PA200-capped proteasomes compared to uncapped ones (Figure 6g, h). These results suggest that the increased turnover efficiency is conserved between Blm10-capped yeast proteasomes and PA200-capped human proteasomes.

## 4. Discussion

The pathological aggregation of αSyn into amyloid fibrils is a central mechanism underlying dopaminergic neuron degeneration in PD. This aggregation process is closely linked to the collapse of proteostasis, the dynamic regulation of protein synthesis, folding, and degradation (Lehtonen et al., 2019). Two key protein degradation systems contribute to cellular proteostasis: the ubiquitin-proteasome system (UPS) and autophagy. Both macroautophagy and chaperone-mediated autophagy are known to facilitate αSyn turnover (Webb et al., 2003; Vogiatzi et al., 2008), yet these processes are dysfunctional in PD patients, with αSyn itself acting as one of the contributors to this inhibition (Fellner et al., 2021). In addition to compromised autophagy, αSyn negatively impacts UPS function. We previously reported that αSyn stabilizes the proteasome assembly chaperone Rpn14, leading to reduced formation of fully active 26S proteasomes (Galka et al., 2024), accompanied by a downregulation in expression of proteasomal subunits (Popova et al., 2021a). In the current study, we extend these findings by exploring on the impact of αSyn on the 20S proteasome and its regulator Blm10. Our results demonstrate that αSyn inhibits the 26S proteasome more severely than the 20S core particle. This is particularly significant as the 20S core particle is catalytically active even in the absence of ATP, unlike the 26S proteasome, which depends on ATP for substrate unfolding and deubiquitination via the 19S regulatory particle (Bard et al., 2018). The activity of the 20S proteasome can be regulated by modulators different than the 19S regulatory particle. This positions the 20S proteasome as a promising alternative degradation route for αSyn. Blm10, the yeast ortholog of human PA200, loosely binds to the 20S α-ring. Both orthologues possess a conserved C-terminal HbYX motif, capable of inducing partial gate opening and enhancing proteolysis (Dange et al., 2011). Prior studies have demonstrated that Blm10- and PA200-capped 20S proteasomes efficiently degrade misfolded disease-associated proteins such as tau and huntingtin (Dange et al., 2011; Aladdin et al., 2020). Here, we demonstrate comprehensively that the Blm10 and PA200-capped proteasomes are key cellular antagonists against the aggregation of amyloid proteins, using the PD related protein αSyn as an example.

This work highlights Blm10-CP as an efficient turn over machine for monomeric αSyn, which is also capable of degrading A11-postive oligomers of αSyn, a thus far unknown function. The performed experiments highlight that the *in vitro* observations translate into an *in vivo* context, through the use of cellular PD models. *In vivo*, *BLM10* overexpression enhances αSyn turnover, reduces aggregation, and restores 20S activity, improving cell viability. Overexpression of *PSME4* also significantly reduces the amount of αSyn inclusions within H4 cells and reduces the steady state levels of αSyn *in vivo*. These novel findings identify 20S proteasomes capped with Blm10 or PA200 as effective αSyn degradation units.

Our data further suggest that *BLM10* overexpression partially rescues 26S proteasome activity, potentially through the formation of hybrid proteasomes, in which the 20S core is capped by the 19S regulatory particle on one side and Blm10 on the other. Although the function of such hybrid proteasomes remains incompletely understood, previous findings (Burris et al., 2021) indicate that Blm10 overexpression alters proteasome composition in favour of non-19S capped complexes. The physiological relevance of this mechanism is highlighted by the conserved function of PA200 in humans. Despite limited sequence similarity, both Blm10 and PA200 function analogously by binding the 20S α-ring and promoting gate opening mediated by the HbYX motif (Yazgili et al., 2022). Importantly, expression of both *BLM10* and *PSME4* declines with age (Chen et al., 2020), potentially contributing to the impaired proteasomal clearance of aggregation prone proteins in age-related diseases such as PD. Moreover, *PSME4* expression is significantly reduced in the blood of PD patients compared to age-matched controls (Yuan et al., 2020), which points to a vulnerability in this proteasome assembly during aging and neurodegeneration. Diminished Blm10/PA200 levels likely impair clearance of αSyn under pathological conditions. PA200 is also involved in the turnover of acetylated histones, a function shared with Blm10 and linked to nuclear proteostasis (Chen et al., 2020). Since αSyn localizes to the nucleus and interferes with histone acetylation (Kontopoulos et al., 2006), nucleosomes may represent a critical site of interaction between αSyn and Blm10/PA200-capped proteasomes. We observed increased Blm10 protein stability upon αSyn expression. However, this stabilization is unlikely to fully compensate for reduced transcriptional levels. Blm10, like PA200, is tightly regulated and present at sub-stoichiometric levels to proteasomes in the cell (Burris et al., 2021). Alternatively, it is possible that the increased stability of Blm10/PA200 could suppress transcription via potential autoregulation. We show a mechanistic link between αSyn-induced autophagy inhibition and Blm10 stabilization. Blm10 is predominantly degraded through autophagy (Burris et al., 2021), whereby αSyn inhibits autophagic flux, specifically autophagosome-vacuole fusion, which has been previously observed in mammalian cells with αSyn pathology (Tang et al., 2021). Impairment of autophagic flux leads to Blm10 stabilization. Notably, this stabilization depends on αSyn phosphorylation at serine 129, further implicating this modification in proteostasis regulation.

Our study provides novel mechanistic insights into αSyn-induced proteasome inhibition. *In vitro* assays revealed that both αSyn oligomers and monomers inhibit 20S activities. This supports prior findings (Thibaudeau et al., 2018), that A11-positive oligomers are potent 20S inhibitors that stabilize a closed proteasome conformation, blocking activation by HbYX-containing regulators. Our observation that monomers also inhibit the 20S core considerably expands this paradigm and suggests a much broader toxic potential of αSyn species. Importantly, Blm10 binding restores activities of the 20S proteasome particles in the presence of both monomeric and oligomeric αSyn, indicating its protective role. Moreover, this effect is conserved when human 20S proteasomes are reconstituted with PA200. Importantly, the observed *in vitro* effect can be recapitulated by overexpression *BLM10* or *PSME4*. These results suggest that Blm10/PA200-capped 20S proteasomes can maintain proteolytic function under αSyn-induced stress, providing an alternate degradation route when the UPS is impaired.

In conclusion, our findings demonstrate that αSyn monomers and oligomers inhibit 20S proteasome function. Most likely this is achieved by stabilizing a closed-gate conformation of the 20S proteasome. Blm10/PA200 binding counteracts this effect, which results in restored proteolytic activity, promoting αSyn clearance. This highlights a unique role for Blm10/PA200-capped 20S proteasomes in maintaining proteostasis under proteotoxic stress. The results provide a new promising perspective, which points to novel therapeutics with potential uses against neurodegenerative diseases including PD as well as other aggregopathies.

## Author contributions

Conceptualization: TTA, BP, GHB, TFO, and ES; funding acquisition: GHB; investigation: TTA, AZ, MM, ZB and BP; supervision: BP, TFO and GHB; writing - original draft: TTA, BP, and GHB; writing - review and editing: TTA, BP, ES, TFO, and GHB.

## Acknowledgments

We thank Jannik Hahn, Ariane Trap, Tessa Jünemann and Masato Ishizaka for their experimental help. This work was supported by the Deutsche Forschungsgemeinschaft (DFG BR1502/21-1 to GB). TFO was supported by DFG SFB1286, Project B8. This work was partly supported by the Göttingen Graduate Centre for Neurosciences, Biophysics and Molecular Biosciences at the Georg-August Universität Göttingen. MM is supported by DAAD Research Grants - Doctoral Programmes in Germany, 57552340.

## Data availability statement

The data that support the findings of this study are available from the corresponding author upon reasonable request.

## References

Aladdin, A., Yao, Y., Yang, C., Kahlert, G., Ghani, M., Király, N., et al. (2020). The Proteasome Activators Blm10/PA200 Enhance the Proteasomal Degradation of N-Terminal Huntingtin. Biomol. 2020, Vol. 10, Page 1581 10, 1581. doi: 10.3390/BIOM10111581

Anderson, J. P., Walker, D. E., Goldstein, J. M., de Laat, R., Banducci, K., Caccavello, R. J., et al. (2006). Phosphorylation of Ser-129 is the dominant pathological modification of alpha-synuclein in familial and sporadic Lewy body disease. J Biol Chem 281, 29739–29752. doi: 10.1074/jbc.M600933200

Bard, J. A. M., Goodall, E. A., Greene, E. R., Jonsson, E., Dong, K. C., and Martin, A. (2018). Structure and Function of the 26S Proteasome. Annu. Rev. Biochem. 87, 697–724. doi: 10.1146/ANNUREV-BIOCHEM-062917-011931/CITE/REFWORKS

Bi, M., Du, X., Jiao, Q., Chen, X., and Jiang, H. (2021). Expanding the role of proteasome homeostasis in Parkinson’s disease: beyond protein breakdown. Cell Death Dis. 12. doi: 10.1038/s41419-021-03441-0

Burris, A., Waite, K. A., Reuter, Z., Ockerhausen, S., and Roelofs, J. (2021). Proteasome activator Blm10 levels and autophagic degradation directly impact the proteasome landscape. doi: 10.1016/j.jbc.2021.100468

Chen, L. Bin, Ma, S., Jiang, T. X., and Qiu, X. B. (2020). Transcriptional upregulation of proteasome activator Blm10 antagonizes cellular aging. Biochem. Biophys. Res. Commun. 532, 211–218. doi: 10.1016/j.bbrc.2020.07.003

Chuah, J. J. Y., Thibaudeau, T. A., and Smith, D. M. (2023). Minimal mechanistic component of HbYX-dependent proteasome activation that reverses impairment by neurodegenerative-associated oligomers. Commun. Biol. 6, 725. doi: 10.1038/s42003-023-05082-9

Dange, T., Smith, D., Noy, T., Rommel, P. C., Jurzitza, L., Cordero, R. J. B., et al. (2011). Blm10 protein promotes proteasomal substrate turnover by an active gating mechanism. J. Biol. Chem. 286, 42830–42839. doi: 10.1074/jbc.M111.300178

Fellner, L., Gabassi, E., Haybaeck, J., and Edenhofer, F. (2021). Autophagy in α-synucleinopathies—an overstrained system. Cells 10, 3143. doi: 10.3390/cells10113143

Galka, D., Ali. T.T., Bast, A., Niederleithinger, M., Gerhardt, E., et al. (2024). Inhibition of 26S proteasome activity by α-synuclein is mediated by the proteasomal chaperone Rpn14/PAAF1. Aging Cell 23, e14128. doi: 10.1111/acel.14128

Gietz, D., St Jean, A., Woods, R. A., and Schiestl, R. H. (1992). Improved method for high efficiency transformation of intact yeast cells. Nucleic Acids Res 20, 1425.

Hipp, M. S., Kasturi, P., and Hartl, F. U. (2019). The proteostasis network and its decline in ageing. Nat. Rev. Mol. Cell Biol. 20, 421–435. doi: 10.1038/s41580-019-0101-y

Hsu, L. J., Sagara, Y., Arroyo, A., Rockenstein, E., Sisk, A., Mallory, M., et al. (2000). α-Synuclein Promotes Mitochondrial Deficit and Oxidative Stress. Am. J. Pathol. 157, 401–410. doi: 10.1016/S0002-9440(10)64553-1

Kleinknecht, A., Popova, B., Lázaro, D. F., Pinho, R., Valerius, O., Outeiro, T. F., et al. (2016). C-Terminal Tyrosine Residue Modifications Modulate the Protective Phosphorylation of Serine 129 of α-Synuclein in a Yeast Model of Parkinson’s Disease. PLOS Genet. 12, e1006098. doi: 10.1371/journal.pgen.1006098

Kontopoulos, E., Parvin, J. D., and Feany, M. B. (2006). Alpha-synuclein acts in the nucleus to inhibit histone acetylation and promote neurotoxicity. Hum Mol Genet 15, 3012–3023. doi: 10.1093/hmg/ddl243

Lázaro, D. F., Rodrigues, E. F., Langohr, R., Shahpasandzadeh, H., Ribeiro, T., Guerreiro, P., et al. (2014). Systematic Comparison of the Effects of Alpha-synuclein Mutations on Its Oligomerization and Aggregation. PLoS Genet. 10, e1004741. doi: 10.1371/journal.pgen.1004741

Lehtonen, Š., Sonninen, T.-M., Wojciechowski, S., Goldsteins, G., and Koistinaho, J. (2019). Dysfunction of Cellular Proteostasis in Parkinson’s Disease. Front. Neurosci. 13, 457. doi: 10.3389/fnins.2019.00457

Li, Y., Tomko, R. J., and Hochstrasser, M. (2015). Proteasomes: Isolation and activity assays. Curr. Protoc. Cell Biol. 2015, 3.43.1–3.43.20. doi: 10.1002/0471143030.cb0343s67

Lindersson, E., Beedholm, R., Højrup, P., Moos, T., Gai, W. P., Hendil, K. B., et al. (2004). Proteasomal Inhibition by α-Synuclein Filaments and Oligomers. J. Biol. Chem. 279, 12924–12934. doi: 10.1074/jbc.M306390200

Maestro-López, M., Cheng, T. C., Muntaner, J., Menéndez, M., Alonso, M., Schweitzer, A., et al. (2025). Structures of the 26S proteasome in complex with the Hsp70 cochaperone Bag1 reveal a novel mechanism of ubiquitin-independent proteasomal degradation. bioRxiv, 2025.01.22.633148. doi: 10.1101/2025.01.22.633148

Mayor, T., Sharon, M., and Glickman, M. H. (2016). Tuning the proteasome to brighten the end of the journey. Am. J. Physiol. Cell Physiol. 311, C793–C804. doi: 10.1152/AJPCELL.00198.2016

McNaught, K. S. P., Belizaire, R., Isacson, O., Jenner, P., and Olanow, C. W. (2003). Altered proteasomal function in sporadic Parkinson’s disease. Exp. Neurol. 179, 38–46. doi: 10.1006/exnr.2002.8050

Mehra, S., Gadhe, L., Bera, R., Sawner, A. S., and Maji, S. K. (2021). Structural and functional insights into α-synuclein fibril polymorphism. Biomolecules 11. doi: 10.3390/biom11101419

Miranda, H. V., Xiang, W., de Oliveira, R. M., Simões, T., Pimentel, J., Klucken, J., et al. (2013). Heat-mediated enrichment of α-synuclein from cells and tissue for assessing post-translational modifications. J. Neurochem. 126, 673–684. doi: 10.1111/jnc.12251

Oueslati, A. (2016). Implication of Alpha-Synuclein Phosphorylation at S129 in Synucleinopathies: What Have We Learned in the Last Decade? J. Parkinsons. Dis. 6, 39–51. doi: 10.3233/JPD-160779

Outeiro, T. F., and Lindquist, S. (2003). Yeast Cells Provide Insight into Alpha-Synuclein Biology and Pathobiology. Science 302, 1772–1775. doi: 10.1126/science.1090439

Petroi, D., Popova, B., Taheri-Talesh, N., Irniger, S., Shahpasandzadeh, H., Zweckstetter, M., et al. (2012). Aggregate clearance of α-synuclein in Saccharomyces cerevisiae depends more on autophagosome and vacuole function than on the proteasome. J. Biol. Chem. 287, 27567–79. doi: 10.1074/jbc.M112.361865

Popova, B., Galka, D., Häffner, N., Wang, D., Schmitt, K., Valerius, O., et al. (2021a). α-Synuclein Decreases the Abundance of Proteasome Subunits and Alters Ubiquitin Conjugates in Yeast. Cells 2021, Vol. 10, Page 2229 10, 2229. doi: 10.3390/CELLS10092229

Popova, B., Kleinknecht, A., and Braus, G. H. (2015). Posttranslational Modifications and Clearing of alpha-Synuclein Aggregates in Yeast. Biomolecules 5, 617–634. doi: 10.3390/biom5020617

Popova, B., Wang, D., Pätz, C., Akkermann, D., Lázaro, D. F., Galka, D., et al. (2021b). DEAD-box RNA helicase Dbp4/DDX10 is an enhancer of α-synuclein toxicity and oligomerization. PLOS Genet. 17, e1009407. doi: 10.1371/journal.pgen.1009407

Sahoo, S., Padhy, A. A., Kumari, V., and Mishra, P. (2022). Role of Ubiquitin–Proteasome and Autophagy-Lysosome Pathways in α-Synuclein Aggregate Clearance. Mol. Neurobiol. 59, 5379–5407. doi: 10.1007/s12035-022-02897-1

Schmidt, M., Haas, W., Crosas, B., Santamaria, P. G., Gygi, S. P., Walz, T., et al. (2005). The HEAT repeat protein Blm10 regulates the yeast proteasome by capping the core particle. Nat. Struct. Mol. Biol. 12, 294–303. doi: 10.1038/nsmb914

Shahpasandzadeh, H., Popova, B., Kleinknecht, A., Fraser, P. E., Outeiro, T. F., and Braus, G. H. (2014). Interplay between sumoylation and phosphorylation for protection against alpha-synuclein inclusions. J Biol Chem 289, 31224–31240. doi: 10.1074/jbc.M114.559237

Spillantini, M. G., Schmidt, M. L., Lee, V. M. Y., Trojanowski, J. Q., Jakes, R., and Goedert, M. (1997). α-Synuclein in Lewy bodies. Nat. 1997 3886645 388, 839–840. doi: 10.1038/42166

Stefanis, L., Emmanouilidou, E., Pantazopoulou, M., Kirik, D., Vekrellis, K., and Tofaris, G. K. (2019). How is alpha-synuclein cleared from the cell? J. Neurochem., jnc.14704. doi: 10.1111/jnc.14704

Tang, Q., Gao, P., Arzberger, T., Höllerhage, M., Herms, J., Höglinger, G., et al. (2021). Alpha-Synuclein defects autophagy by impairing SNAP29-mediated autophagosome-lysosome fusion. Cell Death Dis. 12, 1–16. doi: 10.1038/s41419-021-04138-0

Thibaudeau, T. A., Anderson, R. T., and Smith, D. M. (2018). A common mechanism of proteasome impairment by neurodegenerative disease-associated oligomers. Nat. Commun. 9, 1–14. doi: 10.1038/s41467-018-03509-0

Tofaris, G. K., Layfield, R., and Spillantini, M. G. (2001). α-Synuclein metabolism and aggregation is linked to ubiquitin-independent degradation by the proteasome. FEBS Lett. 509, 22–26. doi: 10.1016/S0014-5793(01)03115-5

Vogiatzi, T., Xilouri, M., Vekrellis, K., and Stefanis, L. (2008). Wild type alpha-synuclein is degraded by chaperone-mediated autophagy and macroautophagy in neuronal cells. J Biol Chem 283, 23542–23556. doi: 10.1074/jbc.M801992200

Webb, J. L., Ravikumar, B., Atkins, J., Skepper, J. N., and Rubinsztein, D. C. (2003). α-synuclein Is Degraded by Both Autophagy and the Proteasome. J. Biol. Chem. 278, 25009–25013. doi: 10.1074/jbc.M300227200

Weberruss, M. H., Savulescu, A. F., Jando, J., Bissinger, T., Harel, A., Glickman, M. H., et al. (2013). Blm10 facilitates nuclear import of proteasome core particles. EMBO J. 32, 2697–2707. doi: 10.1038/emboj.2013.192

Winner, B., Jappelli, R., Maji, S. K., Desplats, P. A., Boyer, L., Aigner, S., et al. (2011). In vivo demonstration that α-synuclein oligomers are toxic. Proc. Natl. Acad. Sci. U. S. A. 108, 4194–4199. doi: 10.1073/pnas.1100976108

Yazgili, A. S., Ebstein, F., and Meiners, S. (2022). The Proteasome Activator PA200/PSME4: An Emerging New Player in Health and Disease. Biomolecules 12. doi: 10.3390/biom12081150

Yuan, Q., Zhang, S., Li, J., Xiao, J., Li, X., Yang, J., et al. (2020). Comprehensive analysis of core genes and key pathways in Parkinson’s disease. Am. J. Transl. Res. 12, 5630.

