## Supplementary information for "Blm10/PA200-activated 20S proteasomes promote α-synuclein degradation and bypass proteasome inhibition in Parkinson’s disease models"

#### Supplementary Tables

##### Table S1. Yeast strains used in this study.

| Name | Description | Source |
|---|---|---|
| W303-1A | MAT*a, ura3–1, trp1–1, leu2–3_112, his3–11, ade2–1, can1–100* | EUROSCARF |
| BY4741 | MAT*a, ura3∆0, his3∆1, leu2∆0, met15∆0* | EUROSCARF |
| *BLM10*-tFT | YMaM330 *BLM10::mCherry::sfGFP* | (Khmelinskii et al., 2012) |
| *BLM10*-tFT ∆*atg7* | YMaM330 *BLM10-tFT*, *Δatg7::KanMX4* | this study |
| *PRE1-3xFLAG* | W303 *ade2-1, ura3-1, his3-11,25, trp1-1, leu2-3,112, can1, pre1::PRE1-3xFLAG-HIS3* | (Saeki et al., 2009) |
| *RPN11-3xFLAG* | W303 *ade2-1, ura3-1, his3-11,15, trp1-1, leu2-3,112, can1, rpn11::RPN11-3xFLAG* | (Saeki et al., 2009) |
| BY4741∆*blm10* | BY4741 *MATa; his3Δ1; leu2Δ0; met15Δ0; ura3Δ0 Δblm10:NatMX4* | Invitrogen deletion collection |
| RH3468 | MAT*α*; *ura3-52; trp1D2; leu2-3_112; his3-11; ade2-1; can1-100 GAL1::SNCA::GFP::URA3* (3 copies) | (Petroi et al., 2012) |
| EGY48 | MATa, *his3, trp1, ura3, LexAop (x6)-LEU2* | (Golemis et al., 1999a) |

##### Table S2. Plasmids used in this study.

| Name | Description | Source |
|---|---|---|
| p426 | *2µ, URA3,* $GAL1_{pr}$, $CYC1_{term}$, *AmpR* | (Mumberg et al., 1994) |
| pME3760 | p426-$GAL1^{pr}$*::SNCA* | (Petroi et al., 2012) |
| pME3763 | p426-$GAL1^{pr}$*::SNCA-GFP* | (Petroi et al., 2012) |

| | | |
|---|---|---|
| pME5320 | p426-*GAL1*$^{pr}$*::SNCA*$^{S129A}$ | (Popova et al., 2021a) |
| pME5570 | p426-*GAL1*$_{pr}$*::SNCA*$^{S129D}$ | this study |
| p425 | *2µ, LEU2, GPD*$_{pr}$*, CYC1*$_{term}$*, AmpR* | (Mumberg et al., 1994) |
| pME5571 | *CEN, LEU2, GPD*$_{pr}$*, CYC1*$_{term}$*, AmpR, mCherryBlm10* | this study |
| pME5572 | p425-*mCherry-BLM10* | this study |
| pME5574 | p426-*3xFLAG-BLM10* | this study |
| pME5575 | p426-*3xFLAG-PA200* | this study |
| pRS315-EGFP-Atg8 | *CEN, LEU2, CYC1*$_{term,}$ *EGFP-ATG8* | (Voigt and Pöggeler, 2013) |
| pME4913 | pET22b-*SNCA* | (Popova et al., 2021c) |
| pEG202 | *2µ, HIS3, ADH*$^{pr}$*, LexA, ADH*$^{term}$ | (Golemis et al., 1999a) |
| pJG4-5 | *2µ, TRP1, GAL1*$^{pr}$*, B42, ADH*$^{term}$ | (Golemis et al., 1999a) |
| pME5525 | pEG202-*SNCA*$^{WT}$ | (Galka et al., 2024) |
| pME5530 | pEG202-*SNCA*$^{S129A}$ | (Galka et al., 2024) |
| pME5573 | pJG4-5-*BLM10* | this study |
| pME5090 | p426-*GAL1*$^{pr}$*::VenusN* | (Popova, et al. 2021) |
| pME5036 | p423-*GAL1*$^{pr}$*::VenusC* | (Popova, et al. 2021) |
| pME5033 | p423-*GAL1::SNCA::VenusC* | (Tenreiro et al., 2016) |
| pME5034 | p426-*GAL1::VenusN::SNCA* | (Tenreiro et al., 2016) |
| pME5517 | p423-*GAL1pr::SNCAS129A::VenusC* | (Galka et al., 2024) |
| pME5518 | p423-*GAL1pr::SNCAS129D::VenusC* | (Galka et al., 2024) |
| pME5577 | p423-*GAL1::BLM10::VenusC* | this study |
| pME5578 | pcDNA 3.1 with 6xHis-PA200 | this study |

**Table S3. Antibodies used in this study.**

| **Name** | **concentration** | **Origin** | **Source** |
|---|---|---|---|
| α-RFP | 1:5000 | rat | Proteintech Group, Rosemont, IL |
| α-GFP | 1:1000 | rat | Proteintech Group, Rosemont, IL |

| | | | |
|---|---|---|---|
| α-FLAG | 1:5000 | mouse | Proteintech Group, Rosemont, IL |
| α-αSyn | 1:2000 | mouse | Becton Dickinson, Franklin Lakes, NJ |
| α-αSyn pS129 | 1:2000 | mouse | WAKO chemicals, Richmond, VA |
| α-A11 | 1:1000 | rabbit | Thermo Fischer, Waltham, MA |
| α-GAPDH | 1:5000 | mouse | Thermo Fischer, Waltham, MA |
| α-rat-HRP | 1:1000 | goat | Thermo Fischer, Waltham, MA |
| α-mouse-HRP | 1:5000 | goat | Thermo Fischer, Waltham, MA |
| α-rabbit-HRP | 1:5000 | goat | Thermo Fischer, Waltham, MA |

#### Supplementary Methods

##### Yeast-Two Hybrid assay

Putative protein-protein interaction were analyzed using yeast-two-hybrid assay as previously described (Golemis et al., 1999b). Proteins are genetically fused to either a DNA binding domain or an activation domain of a transcriptional activator. α-synuclein and its variants S129A and S129D were fused to the activation domain, while Blm10 was attached to the DNA binding domain of the bacterial repressor protein LexA. Constructs were co-transformed into the yeast strain EGY48 and interaction was analyzed using selective SC-His-Trp-Leu plates supplemented with 2% galactose or 2% glucose as a negative control.

##### Bimolecular fluorescence complementation assay

Potential interactions between Blm10 and αSyn or its variants were analyzed using bimolecular fluorescence complementation (BiFC) assay (Hu et al., 2002). The Venus fluorophore was split into C- and N-terminal fragments, which were genetically fused to the proteins of interest. Blm10 was fused with the N-terminal fragment, whereas αSyn, S129A, and S129D were fused to the C-terminal fragment. The constructs were co-transformed into the BY4741 yeast strain and expressed, followed by fluorescence microscopy. Protein-protein interaction was indicated by a reconstituted Venus signal detectable in the GFP channel.

#### Nitrogen Starvation

Transformed yeast cells were grown overnight in selective SC medium supplemented with 2% raffinose. The cultures were then diluted to an $OD_{600}$ of 0.3 and re-inoculated in SC medium containing 2% galactose to induce *GAL1*-driven expression of αSyn and its mutants for 6 h. After induction, samples were collected for microscopic analysis and protein extraction. The remaining cells were washed twice and transferred to nitrogen-deficient SC medium containing 0.2% galactose. After 16 h of starvation, cells were harvested for protein extraction and microscopic analysis.

#### References


Galka, D., Ali, T. T., Bast, A., Niederleithinger, M., Gerhardt, E., Motosugi, R., et al. (2024). Inhibition of 26S proteasome activity by α-synuclein is mediated by the proteasomal chaperone Rpn14/PAAF1. *Aging Cell* 23. doi:10.1111/acel.14128.

Golemis, E. A., Serebriiskii, I., and Law, S. F. (1999a). The yeast two-hybrid system: criteria for detecting physiologically significant protein-protein interactions. *Curr. Issues Mol. Biol.* 1, 31–45. doi:10.21775/cimb.001.031.

Golemis, E. A., Serebriiskii, I., and Law, S. F. (1999b). The yeast two-hybrid system: criteria for detecting physiologically significant protein-protein interactions. *Curr. Issues Mol. Biol.* 1, 31–45. doi:10.21775/CIMB.001.031.

Hu, C. D., Chinenov, Y., and Kerppola, T. K. (2002). Visualization of Interactions among bZIP and Rel Family Proteins in Living Cells Using Bimolecular Fluorescence Complementation. *Mol. Cell* 9, 789–798. doi:10.1016/S1097-2765(02)00496-3.

Khmelinskii, A., Keller, P. J., Bartosik, A., Meurer, M., Barry, J. D., Mardin, B. R., et al. (2012). Tandem fluorescent protein timers for in vivo analysis of protein dynamics. *Nat. Biotechnol.* 30, 708–714. doi:10.1038/nbt.2281.

Mumberg, D., Muller, R., Funk, M., Müller, R., and Funk, M. (1994). Regulatable promoters of Saccharomyces cerevisiae: comparison of transcriptional activity and their use for heterologous expression. *Nucleic Acids Res.* 22, 5767–8.

Petroi, D., Popova, B., Taheri-Talesh, N., Irniger, S., Shahpasandzadeh, H., Zweckstetter, M., et al. (2012). Aggregate clearance of alpha-synuclein in *Saccharomyces cerevisiae* depends more on autophagosome and vacuole function than on the proteasome. *J Biol Chem* 287, 27567–27579. doi:M112.361865 [pii]10.1074/jbc.M112.361865.

Popova, B., Galka, D., Häffner, N., Wang, D., Schmitt, K., Valerius, O., et al. (2021a). α-Synuclein Decreases the Abundance of Proteasome Subunits and Alters Ubiquitin Conjugates in Yeast. *Cells 2021, Vol. 10, Page 2229* 10, 2229. doi:10.3390/CELLS10092229.

Popova, B., Wang, D., Pätz, C., Akkermann, D., Lázaro, D. F., Galka, D., et al. (2021b). DEAD-box RNA helicase Dbp4/DDX10 is an enhancer of α-synuclein toxicity and oligomerization. *PLOS Genet.* 17, e1009407. doi:10.1371/journal.pgen.1009407.

Popova, B., Wang, D., Rajavel, A., Dhamotharan, K., Lázaro, D. F., Gerke, J., et al. (2021c). Identification of Two Novel Peptides That Inhibit α-Synuclein Toxicity and Aggregation. *Front. Mol. Neurosci.* 14, 54. doi:10.3389/FNMOL.2021.659926.

Saeki, Y., Toh-e, A., Kudo, T., Kawamura, H., and Tanaka, K. (2009). Multiple Proteasome-Interacting Proteins Assist the Assembly of the Yeast 19S Regulatory Particle. *Cell* 137, 900–913. doi:10.1016/J.CELL.2009.05.005.

Tenreiro, S., Rosado-Ramos, R., Gerhardt, E., Favretto, F., Magalhães, F., Popova, B., et al. (2016). Yeast reveals similar molecular mechanisms underlying alpha- and beta-synuclein toxicity. *Hum. Mol. Genet.* 25, 275–290. doi:10.1093/hmg/ddv470.

Voigt, O., and Pöggeler, S. (2013). Autophagy genes Smatg8 and Smatg4 are required for fruiting-body development, vegetative growth and ascospore germination in the filamentous ascomycete Sordaria macrospora. *Autophagy* 9, 33–49. doi:10.4161/AUTO.22398.

### Supplementary Figures

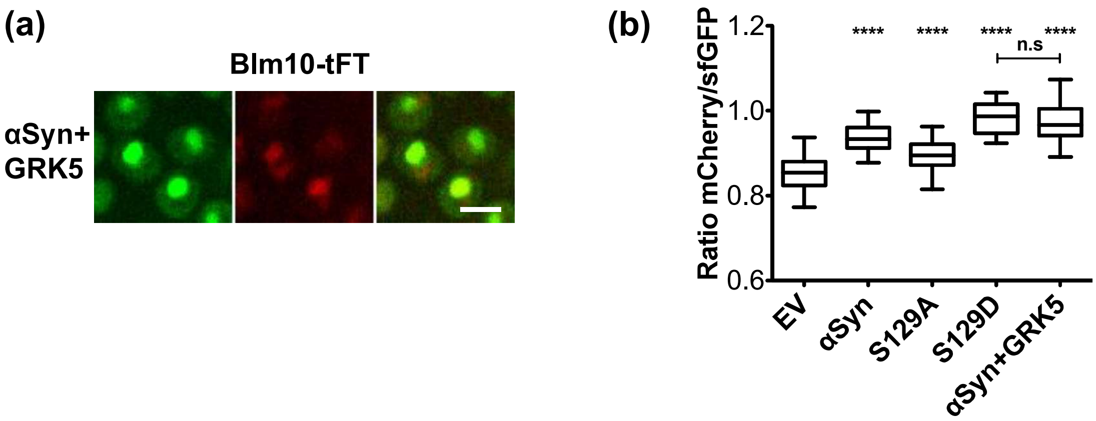


**Figure S1. Hyperphosphorylation of αSyn stabilizes Blm10 similarly to the phospho-mimicking mutant S129D. (a)** Tandem Fluorescent Timer (tFT) analysis of Blm10 stability by fluorescence microscopy. Cells express *GAL1*-driven αSyn and constitutively expressed *GRK5*. Images were acquired after 6 h of αSyn induction. Scale bar = 5 µm. **(b)** Quantification of the mCherry to sfGFP fluorescence ratio from microscopy images, compared to samples shown in Figure 1. Fluorescence ratios were calculated for single cells. Statistical significance was determined by one-way ANOVA with Dunnett's post hoc test (*****p* < 0.0001; n.s *p* > 0.05; n = 50) in comparison to control empty vector (EV).

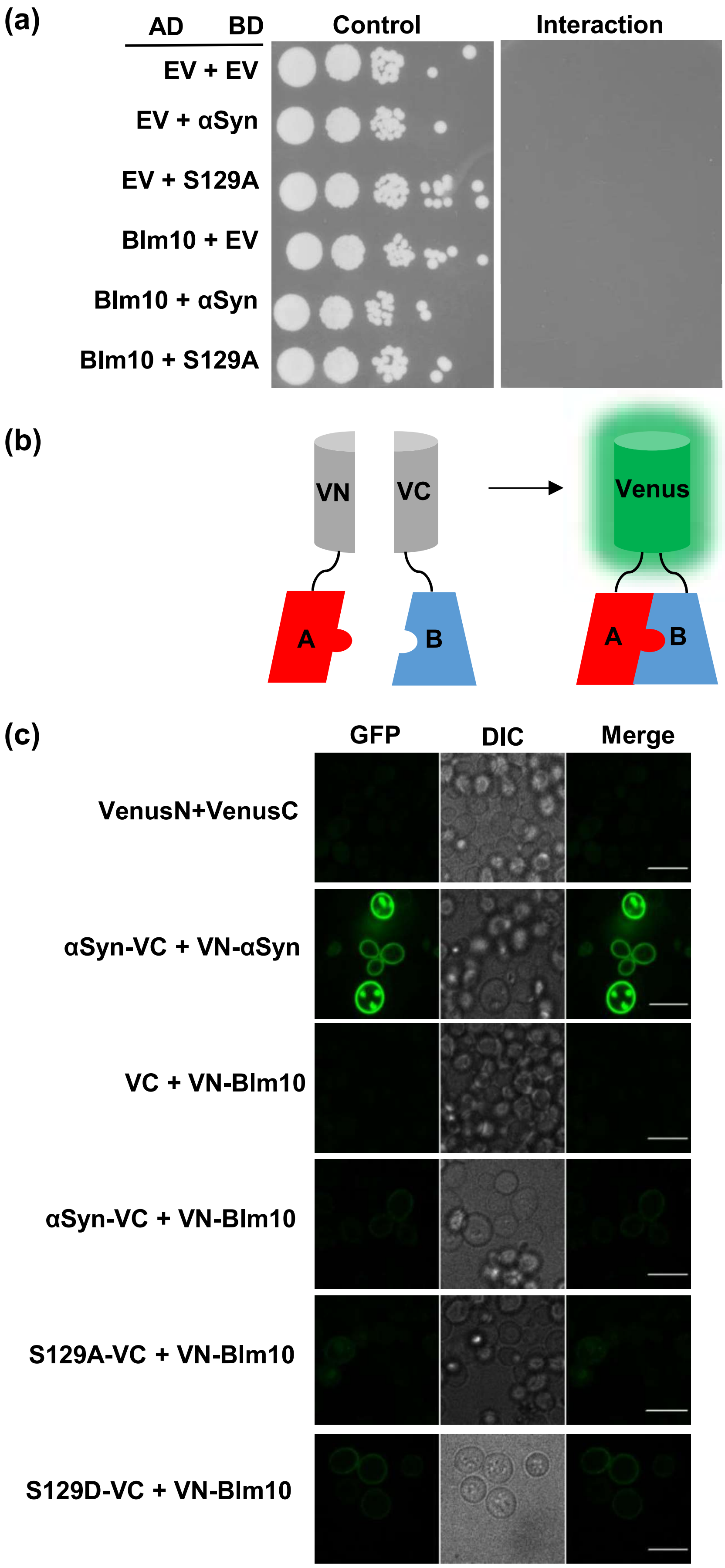
(a)
AD BD
Control
Interaction
EV + EV
EV + αSyn
EV + S129A
Blm10 + EV
Blm10 + αSyn
Blm10 + S129A
(b)
VN
VC
Venus
A
B
(c)
GFP
DIC
Merge
VenusN+VenusC
αSyn-VC + VN-αSyn
VC + VN-Blm10
αSyn-VC + VN-Blm10
S129A-VC + VN-Blm10
S129D-VC + VN-Blm10

**Figure S2. Blm10 and αSyn do not interact physically.** Yeast-two-hybrid assay using yeast cells transformed with plasmids encoding proteins fused to either a transcriptional activation domain (AD) or a DNA-binding domain (BD). The B42 activation domain-fused protein (prey) is under *GAL1* promotor control, while the LexA-DNA BD (bait) is under constitutive *ADH* promotor control. *LEU2* was used as a reporter gene. The control plate contains leucine and is used as control for equal dilution. The interaction plate lacks leucine which permits growth only to cells where AD and BD fusion proteins interact, thus enabling expression of *LEU2* gene. (b) Schematic representation of Bimolecular fluorescence complementation assay (BiFC). Proteins of interest are genetically fused to the C- and N-terminal domains of the improved YFP version Venus (VN and VC). Upon interaction of the proteins, Venus fluorescence is reconstituted. (c) Fluorescence microscopy images of BiFC analyzing the potential interaction between Blm10 and αSyn after 6 h of induction. VenusN + VenusC serves as a negative control and αSyn-VC + αSyn-VN as positive control. Scale bar = 5 µm.

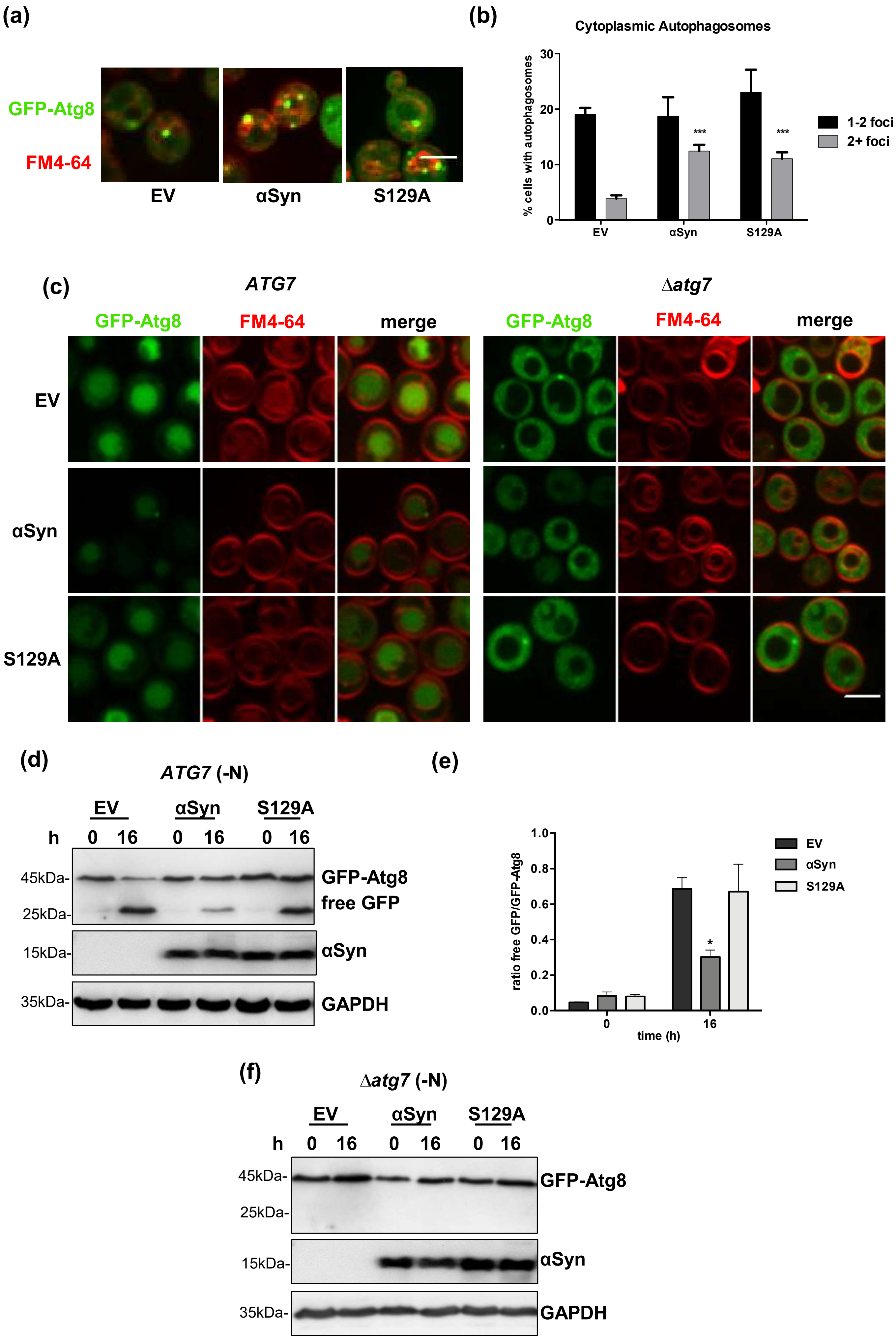

(a)
GFP-Atg8
FM4-64
EV
αSyn
S129A
(b)
Cytoplasmic Autophagosomes
% cells with autophagosomes
30
20
10
0
***
***
1-2 foci
2+ foci
EV
αSyn
S129A
(c)
ATG7
Δatg7
GFP-Atg8
FM4-64
merge
EV
αSyn
S129A
(d)
ATG7 (-N)
EV
αSyn
S129A
h 0 16 0 16 0 16
45kDa-
25kDa-
GFP-Atg8
free GFP
15kDa-
αSyn
35kDa-
GAPDH
(e)
ratio free GFP/GFP-Atg8
1.0
0.8
0.6
0.4
0.2
0.0
*
0
16
time (h)
EV
αSyn
S129A
(f)
Δatg7 (-N)
EV
αSyn
S129A
h 0 16 0 16 0 16
45kDa-
25kDa-
GFP-Atg8
15kDa-
αSyn
35kDa-
GAPDH

**Figure S3. αSyn inhibits autophagy. (a)** Fluorescence microcopy images of yeast cells expressing GFP-tagged Atg8 to monitor autophagosomes. FM4-64 dye was used to label the vacuolar membrane. Cells were imaged after 6 h αSyn expression. Scale bar = 5 µm. **(b)** Quantification of cells exhibiting 1-2 or more autophagosomes. Statistical significance was determined by one-way ANOVA with Dunnett's post hoc test (***$p < 0.001$; n = 3) in comparison to the empty vector (EV) control. **(c)** Fluorescence microscopy images of cells after 16 h of nitrogen starvation to induce autophagy. αSyn expression was induced for 6 h before induction of autophagy. GFP-Atg8 was used as an autophagy marker, with vacuolar GFP signal indicating autophagic flux. FM4-64 staining was used to visualize the vacuolar membrane. The Atg7 deletion mutant, which impairs autophagy, was used as a control. Scale bar = 5 µm. **(d)** Immunoblot analysis of yeast cells expressing GFP-Atg8 before (0 h) and after 16 h of nitrogen starvation. Autophagic flux was evaluated by detecting the release of free GFP from GFP-Atg8. αSyn expression was induced for 6 h prior to induction of autophagy. αSyn expression was conformed with αSyn antibody and GAPDH was used as a loading control. **(e)** Densitometric quantification of the ratio of free GFP to GFP-Atg8 based on scanned immunoblots. Statistical significance was determined by one-way ANOVA with Dunnett's post hoc test (*$p < 0.05$; n = 3) in comparison to the empty vector (EV) control. **(f)** Immunoblot analysis of yeast cells deficient in autophagy expressing GFP-Atg8, before and after 16 h of nitrogen starvation. Analysis was performed as described in (d).

**250 µM 20S + 250 µM Blm10** **250 µM 20S + 500 µM Blm10**

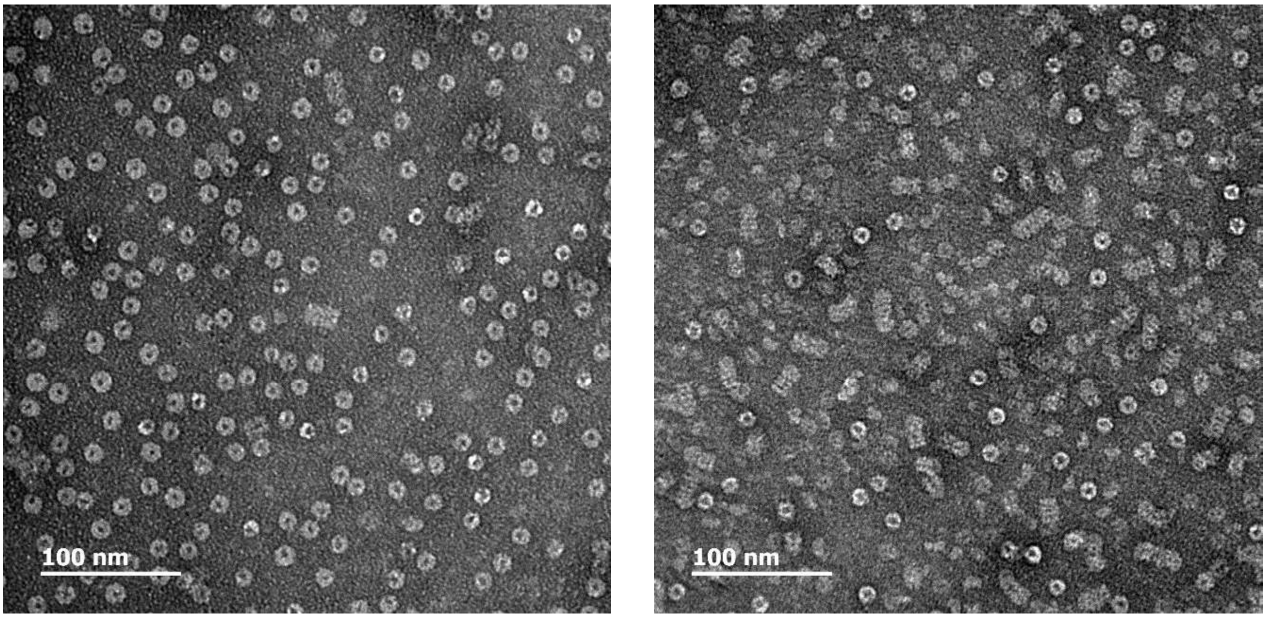


**Figure S4. Reconstitution of 20S proteasomes with Blm10.** Transmission electron microscopy (TEM) images of 20S proteasomes reconstituted with varying concentrations of Blm10. Reconstitution was performed by incubating purified Blm10 with purified 20S proteasome at indicated concentration for 30 min at 30 °C. Samples were applied to CF200-Cu grids, stained with 2% uranyl acetate, and imaged.

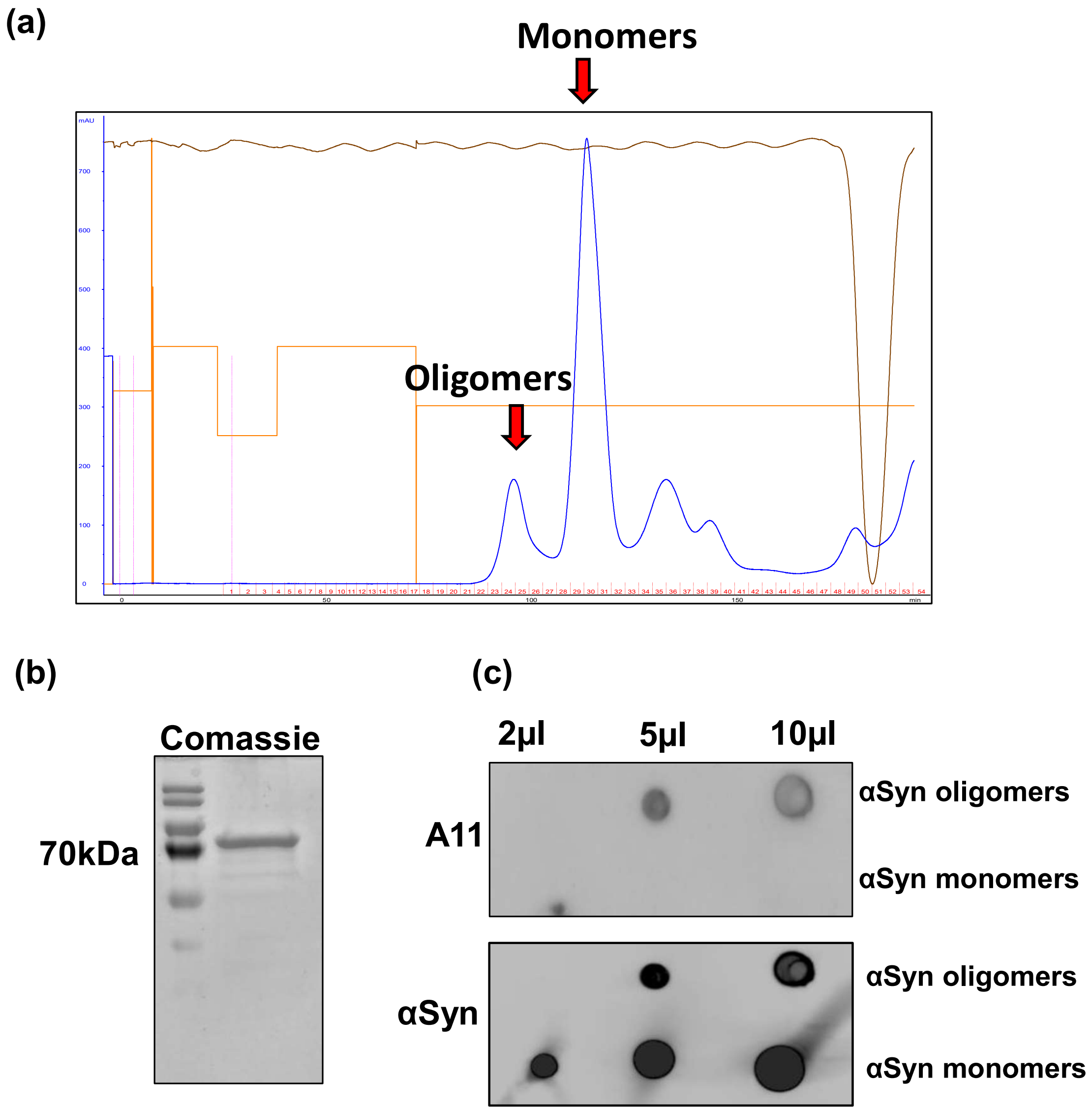


**Figure S5. Purification of αSyn oligomers for activity assays. (a)** Chromatogram of size exclusion chromatography (SEC) of αSyn purification from *E.coli*. Blue curve represents absorption at 280 nm. Two major peaks were defined, the first corresponds to αSyn oligomers, and the second to αSyn monomers. **(b)** Comassie staining of SDS-PAGE gel of oligomer fraction following SEC. Heat and SDS-stable αSyn oligomers were detected at 70 kDa mark. **(c)** Dot blot analysis of αSyn oligomeric and monomeric fractions obtained from SEC. The A11 antibody, specific for oligomeric amyloidogenic species, was used along with αSyn antibody as a control.

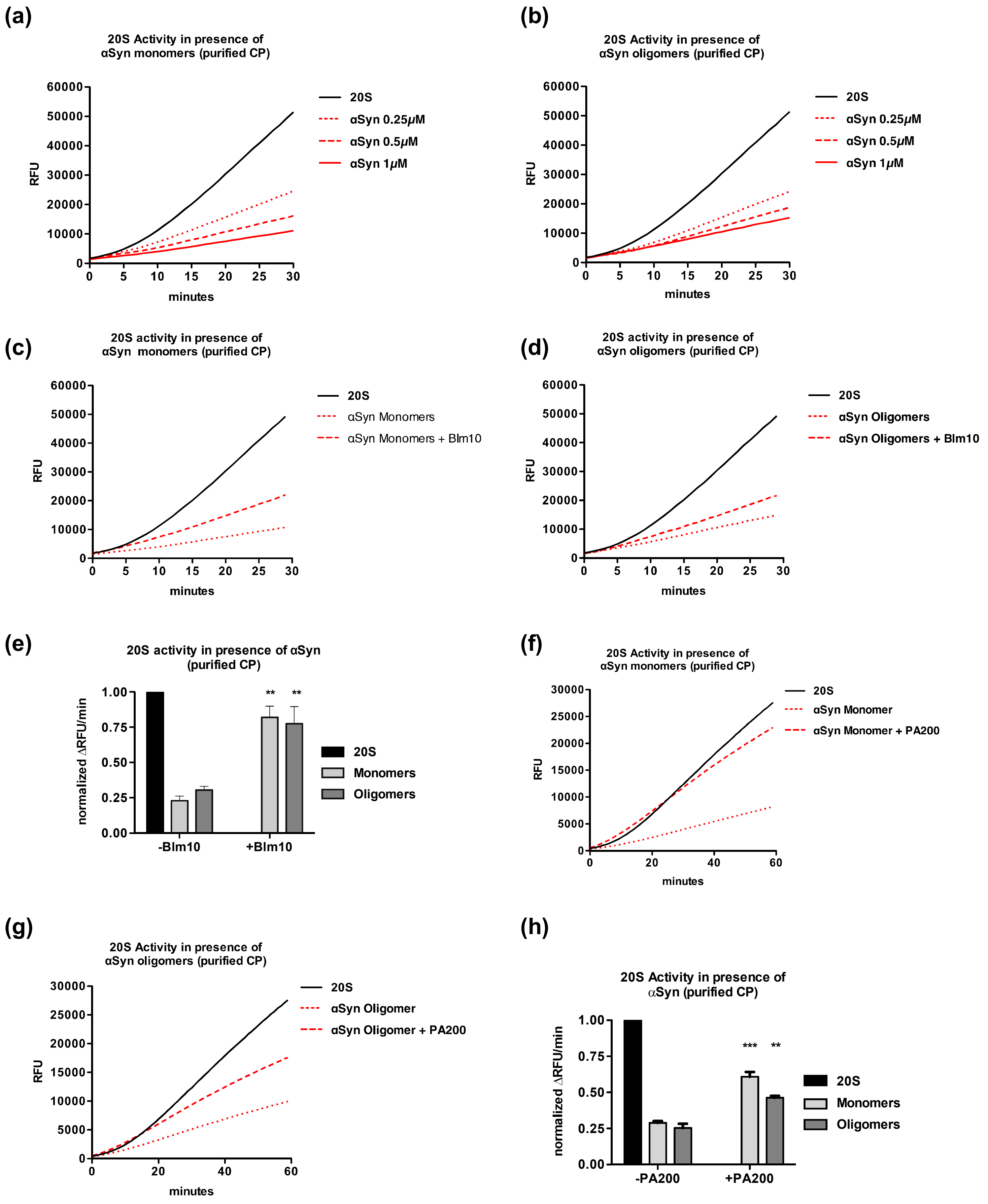
(a)
20S Activity in presence of
αSyn monomers (purified CP)
RFU
minutes
20S
αSyn 0.25μM
αSyn 0.5μM
αSyn 1μM
(b)
20S Activity in presence of
αSyn oligomers (purified CP)
RFU
minutes
20S
αSyn 0.25μM
αSyn 0.5μM
αSyn 1μM
(c)
20S activity in presence of
αSyn monomers (purified CP)
RFU
minutes
20S
αSyn Monomers
αSyn Monomers + Blm10
(d)
20S activity in presence of
αSyn oligomers (purified CP)
RFU
minutes
20S
αSyn Oligomers
αSyn Oligomers + Blm10
(e)
20S activity in presence of αSyn
(purified CP)
normalized ΔRFU/min
**
**
-Blm10
+Blm10
20S
Monomers
Oligomers
(f)
20S Activity in presence of
αSyn monomers (purified CP)
RFU
minutes
20S
αSyn Monomer
αSyn Monomer + PA200
(g)
20S Activity in presence of
αSyn oligomers (purified CP)
RFU
minutes
20S
αSyn Oligomer
αSyn Oligomer + PA200
(h)
20S Activity in presence of
αSyn (purified CP)
normalized ΔRFU/min
***
**
-PA200
+PA200
20S
Monomers
Oligomers

**Figure S6. αSyn oligomers and monomers inhibit 20S proteasome activity. (a-b)** SUC-LLVY-AMC activity assays of purified 20S proteasomes in the absence and presence of increasing concentrations of αSyn monomers (a) or oligomers (b). Graphs represent the mean of 3 replicates. **(c-d)** SUC-LLVY-AMC activity assays of 20S proteasomes and reconstituted 20S+Blm10 complexes in the presence of 1 μM αSyn monomers (c) or oligomers (d). Activity of 20S proteasomes alone (without Blm10 and αSyn) served as control. Data represent the mean of 3 replicates. **(e)** Mean change in fluorescence per minute calculated from (c) and (d). Statistical significance was assessed using one-way ANOVA with Dunnett's post-hoc test (**$p< 0.01$; n=3). **(f-g)** SUC-LLVY-AMC activity assays of 20S proteasomes and reconstituted 20S+PA200 complexes in the presence of 1 μM αSyn monomers (f) or oligomers (g). Activity of 20S proteasomes alone (without PA200 and αSyn) served as control. Data represent the mean of 3 replicates. **(h)** Mean change in fluorescence per minute of human 20S proteasomes with and without reconstitution with PA200 in the presence of αSyn monomers and oligomers. Statistical significance was assessed using one-way ANOVA with Dunnett's post-hoc test (***$p< 0.001$; n=3).
